# Neuronal PAS domain 1 identifies a major subpopulation of wakefulness-promoting GABAergic neurons in basal forebrain

**DOI:** 10.1101/2023.11.09.566065

**Authors:** Timothy A. Troppoli, Chun Yang, Fumi Katsuki, David S. Uygun, Ilyan Lin, David Aguilar, Tristan Spratt, Radhika Basheer, James M. McNally, C. Savio Chan, James T. McKenna, Ritchie E. Brown

## Abstract

Here we describe a novel group of basal forebrain (BF) neurons expressing neuronal PAS domain 1 (Npas1), a developmental transcription factor linked to neuropsychiatric disorders. Immunohistochemical staining in Npas1-cre-2A-TdTomato mice revealed BF Npas1^+^ neurons are distinct from well-studied parvalbumin or cholinergic neurons. Npas1 staining in GAD67-GFP knock-in mice confirmed that the vast majority of Npas1^+^ neurons are GABAergic, with minimal colocalization with glutamatergic neurons in vGlut1-cre-tdTomato or vGlut2-cre-tdTomato mice. The density of Npas1^+^ neurons was high, 5-6 times that of neighboring cholinergic, parvalbumin or glutamatergic neurons. Anterograde tracing identified prominent projections of BF Npas1^+^ neurons to brain regions involved in sleep-wake control, motivated behaviors and olfaction such as the lateral hypothalamus, lateral habenula, nucleus accumbens shell, ventral tegmental area and olfactory bulb. Chemogenetic activation of BF Npas1^+^ neurons in the light (inactive) period increased the amount of wakefulness and the latency to sleep for 2-3 hr, due to an increase in long wake bouts and short NREM sleep bouts. Non-REM slow-wave (0-1.5 Hz) and sigma (9-15 Hz) power, as well as sleep spindle density, amplitude and duration, were reduced, reminiscent of findings in several neuropsychiatric disorders. Together with previous findings implicating BF Npas1^+^ neurons in stress responsiveness, the anatomical projections of BF Npas1^+^ neurons and the effect of activating them suggest a possible role for BF Npas1^+^ neurons in motivationally-driven wakefulness and stress-induced insomnia. Identification of this major subpopulation of BF GABAergic neurons will facilitate studies of their role in sleep disorders, dementia and other neuropsychiatric conditions involving BF.

**SIGNIFICANCE STATEMENT:** We characterize a group of basal forebrain (BF) neurons in the mouse expressing neuronal PAS domain 1 (Npas1), a developmental transcription factor linked to neuropsychiatric disorders. BF Npas1^+^ neurons are a major subset of GABAergic neurons distinct and more numerous than cholinergic, parvalbumin or glutamate neurons. BF Npas1^+^ neurons target brain areas involved in arousal, motivation and olfaction. Activation of BF Npas1^+^ neurons in the light (inactive) period increased wakefulness and the latency to sleep due to increased long wake bouts. Non-REM sleep slow waves and spindles were reduced reminiscent of findings in several neuropsychiatric disorders. Identification of this major subpopulation of BF GABAergic wake-promoting neurons will allow studies of their role in insomnia, dementia and other conditions involving BF.

## INTRODUCTION

Sleep is one of the three pillars of health alongside diet and exercise, and disrupted sleep has been linked to a wide variety of physical and neuropsychiatric consequences (Brown et al., 2022). Sleep disturbances are common among the general public, with higher prevalence among current and former military personnel, the elderly, and those suffering with psychiatric disorders, neurodegenerative disease, or traumatic brain injury (Brown et al., 2022). Precise delineation of the brain circuits which control sleep and wakefulness is essential for the development of new treatments but is hampered by the lack of markers to identify the key neuronal subpopulations in evolutionarily conserved brain regions.

Different subgroups of neurons releasing the main inhibitory neurotransmitter in the brain, gamma-aminobutyric acid (GABA), are involved in a wide variety of different vigilance state regulatory roles, including the promotion of NREM and REM sleep, wakefulness, and the cortical oscillations which are typical of these states (Brown and McKenna, 2015). The wide distribution of GABAergic neurons makes it challenging to find markers which identify key subpopulations, thus hindering targeted experiments to determine their functionality. An approach that has been productive in defining subpopulations of cerebral cortex GABAergic interneurons (Lim et al., 2018) and subpopulations of GABAergic projection neurons in the basal ganglia (Hernandez et al., 2015; Abecassis et al., 2020; Cui et al., 2021) is the use of developmental transcription factors to serve as markers for specific neuronal populations. This approach has also been fruitful in identifying novel sleep-wake populations in the hypothalamus (Liu et al., 2017) and brainstem (Kirjavainen et al., 2022), but has so far not been applied to the basal forebrain (BF), a key brain region involved in the control of sleep-wake behavior, cortical activation, attention, and motivation (Detari et al., 1999; Zaborszky and Duque, 2003; Brown et al., 2012; Brown and McKenna, 2015; Lin et al., 2015; Morais-Silva et al., 2023) which degenerates in dementia patients (Grothe et al., 2014; Pereira et al., 2020) and is dysregulated in various psychiatric disorders (McNally et al., 2021; Morais-Silva et al., 2023; Soares-Cunha & Heinsbroek 2023).

Previous chemogenetic experiments showed that activation of all GABAergic neurons in the BF leads to a large increase in the amount of wakefulness (Anaclet et al., 2015; Li et al., 2021), whereas chemogenetic activation of cholinergic or glutamatergic BF neurons has less pronounced effects on sleep-wake behavior and cortical EEG (Anaclet et al., 2015). However, BF GABAergic neurons are heterogeneous in their neurochemical phenotype and properties (McKenna et al., 2013; Brown and McKenna, 2015; Yang et al., 2017). To date, only those GABAergic neurons which contain the calcium-binding protein parvalbumin (PV) or the neuropeptide somatostatin (SST) have been analyzed in detail. Optogenetic activation of BF PV^+^ neurons or their projections to the thalamic reticular nucleus (TRN) enhances cortical gamma oscillations and leads to rapid arousals from sleep (Kim et al., 2015; Thankachan et al., 2019; McKenna et al., 2020; McNally et al., 2021). However, arousals elicited by optogenetic activation of BF PV^+^ neurons are brief and only slightly increase the amount of wakefulness (Xu et al., 2015; Thankachan et al., 2019; McKenna et al., 2020). In contrast, chemogenetic activation of BF SST neurons has little effect on sleep-wake states or cortical EEG (Anaclet et al., 2018), and strong optogenetic activation of BF SST neurons is sleep-promoting and inhibits cortical activation (Xu et al., 2015; Espinosa et al., 2019). Thus, the strong increase in wakefulness elicited by activation of all BF GABAergic neurons cannot be explained by activation of BF PV^+^ or SOM^+^ neurons, and a marker for the wake-promoting GABAergic BF population remains to be uncovered.

Neuronal Per-Arnt-Sim domain 1 (Npas1) is a basic helix-loop-helix class transcription factor (Zhou et al., 1997) which has been linked by genetic and molecular studies to neuropsychiatric disorders (Erbel-Sieler et al., 2004; Michaelson et al., 2017). Npas1 is expressed in the precursors of cortical interneurons derived from the caudal ganglionic eminence as well as portions of the medial and lateral ganglionic eminences which generate basal ganglia GABAergic projection neurons (Cobos et al., 2006; Flandin et al., 2010; Nobrega-Periera et al., 2010; Stanco et al., 2014). In the globus pallidus pars externa (GPe), Npas1^+^ GABAergic projection neurons represent a distinct functional class of neurons from PV^+^ neurons (Hernandez et al., 2015), suggesting that Npas1 might represent a marker for non-PV^+^ GABAergic neurons in the BF. Npas1^+^ neurons have previously been reported within the BF region (Erbel-Sieler et al., 2004; Hernandez et al., 2015; Morais-Silva et al., 2023) but their neurochemical phenotype is unclear and their role in sleep-wake control has not been studied.

Here we show that Npas1 identifies a major subpopulation of BF GABAergic neurons, distinct from and considerably more dense than previously characterized PV^+^ neurons, cholinergic or glutamatergic neurons. BF Npas1^+^ neurons project to brain regions involved in sleep-wake control, motivated behavior, and olfaction. When activated chemogenetically, BF Npas1^+^ neurons promote wakefulness and disrupt NREM sleep and cortical oscillations.

## RESULTS

### Many small or medium-sized Npas1-tdTomato^+^ neurons are present in the BF

To study the distribution and neurochemical phenotype of BF Npas1^+^ neurons, we used previously validated Npas1-cre-2A-tdTomato mice (Hernandez et al., 2015) together with immunohistochemical (IHC) staining. The fluorescent marker tdTomato is expressed at low levels in the brains of Npas1-cre-2A-tdTomato mice (Hernandez et al., 2015). Thus, to comprehensively visualize Npas1-tdTomato^+^ neurons, tdTomato fluorescence was amplified using a red fluorescent protein (RFP) IHC stain. Following staining, dense clusters of tdTomato^+^ neurons were observed in all three main regions of the intermediate BF, the substantia innominata/ventral pallidum (SI/VP), horizontal limb of the diagonal band (HDB) and magnocellular preoptic nucleus (MCPO) (**Fig. 1A-C**). Quantification of the densities of Npas1-tdTomato^+^ neurons in these three regions, across rostral, medial, and caudal BF coronal sections revealed that the density was significantly higher in ventral HDB (279.5 ± 24.8 neurons/mm^2^/section, n=3 mice) and MCPO subregions (274.4 ± 24.6 neurons/mm^2^/section) than the in dorsal SI/VP (149.7 ± 9.3 neurons/mm^2^/section; p<0.05, ordinary one-way ANOVA) (**Fig. 1D**). Dorsal to these regions, a lower but still substantial density of Npas1^+^ neurons was found in the bed nucleus of the stria terminalis (BNST; 105.81 ± 7.85 neurons/mm^2^/section). Medially, Npas1-tdTomato^+^ neurons were also located in the lateral and medial preoptic areas (120.82 ± 19.02 and 105.64 ± 13.88 neurons/mm^2^/section respectively). Only a few Npas1-tdTomato^+^ neurons (typically <5/section) were observed in the sleep-promoting ventrolateral preoptic nucleus or median preoptic nucleus. Laterally, scattered interneurons were located in the piriform cortex, as well as in other neocortical regions. Comparing the density of Npas1-tdTomato^+^ neurons with our previously described density measures for other neuronal phenotypes in the BF, it was clear that Npas1^+^ neurons are a major subpopulation (**Fig. 1E**). For instance, in the MCPO region, the density of Npas1^+^ neurons was 48.2% of that for all GABAergic neurons (GAD67+ neurons in GAD67-GFP knock-in mice; McKenna et al., 2013). In this same subregion, the density of Npas1^+^ neurons was ∼5 times that of PV^+^ neurons (McKenna et al., 2013), 5.8 times that of cholinergic neurons (ChAT^+^; Yang et al., 2014) and 4.8 times the density of vesicular glutamate transporter subtype 2 (vGlut2^+^) neurons (McKenna et al., 2021).

**Figure 1.**
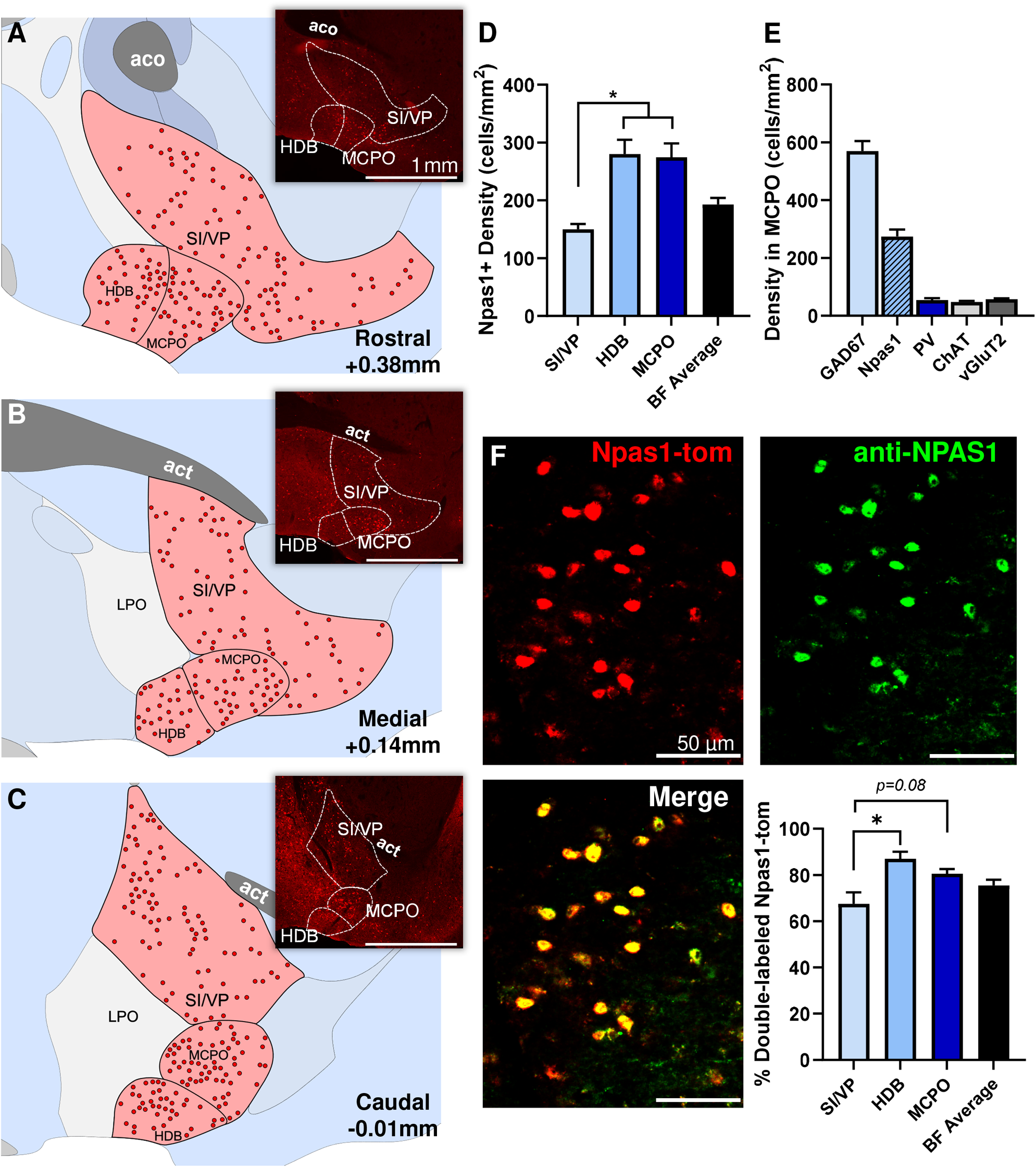
Neurons expressing the transcription factor neuronal PAS domain 1 (Npas1) comprise a major cell population of the basal forebrain (BF). **(A-C)** Npas1^+^ neurons are abundant within the intermediate BF and its major subregions, including the substantia innominata and ventral pallidum (SI/VP), horizontal diagonal band (HDB) and magnocellular preoptic nucleus (MCPO), depicted across **(A)** rostral (+0.38 mm from bregma), **(B)** medial (+0.14 mm), and **(C)** caudal (−0.01 mm) coronal sections. Insets show low power images of Npas1-tdTomato^+^ neurons stained with an anti-red fluorescent protein antibody to enhance endogenous tdTomato fluorescence. Abbreviations: aco, anterior commissure, olfactory limb; act, anterior commissure, temporal limb; LPO, lateral preoptic area. **(D)** Npas1^+^ neurons are denser in ventral (HDB, MCPO) BF subregions compared to SI/VP (n=3 animals; p<0.05, one-way ANOVA, F_(2,6)_ = 12.76, p=0.0069). **(E)** Within the MCPO where GABAergic cell density is highest (McKenna et al., 2013), Npas1^+^ cell density is 48.2% of that of all GABAergic cells. Npas1^+^ cell density is 5-6 times higher than other major cell subpopulations, such as parvalbumin (PV), choline acetyltransferase (ChAT) and vesicular glutamate transporter 2-(vGlut2) expressing neurons (Densities previously published in McKenna et al., 2013, 2021; Yang et al., 2014). **(F)** Although Npas1 is a transcription factor active early in development, the majority (75.5% ± 2.47%, n=3 animals) of BF Npas1-tdTomato neurons (anti-RFP, red) in adult mice continue to express Npas1 as quantified through colocalization with an anti-NPAS1 nuclear stain (green). Colocalization was significantly higher in ventral subregions (HDB, MCPO) compared to the SI/VP (67.5% ± 5.0%; p<0.05, one-way ANOVA, F_(2,6)_ = 7.83, p=0.021). p<0.05*.

Measurements of the long-axis diameter revealed that BF Npas1-tdTomato^+^ neurons were generally small-(10-15 μm) or medium-sized neurons (15-20 μm) (**Supplemental Figure 1A**). Only 3 of 270 Npas1-tdTomato^+^ neurons evaluated had a long-axis diameter >20 μm (**Supplemental Figure 1B**). Overall, BF Npas1-tdTomato^+^ neurons had a mean long-axis diameter of 13.2 ± 0.4 µm (270 neurons in 3 mice). Npas1^+^ neurons were slightly larger in the SI/VP (14.17 ± 0.35 µm) when compared to their counterparts in the HDB (12.63 ± 0.55 µm; p=0.0010) and MCPO (12.70 ± 0.39 µm; p=0.0012, ordinary one-way ANOVA, **Supplemental Figure 1C**). BF Npas1-tdTomato^+^ neurons were smaller than BF PV^+^ neurons (mean long-axis diameter 20.2 µm; McKenna et al., 2013) or cholinergic neurons (24.4 µm; McKenna et al., 2013) but similar in size to all GAD67^+^ neurons (15.9 µm; McKenna et al., 2013, including large PV^+^ neurons) and vGlut2^+^ neurons (13.4 µm; McKenna et al., 2021)

Npas1 is a transcription factor which is active early in development in the ganglionic eminence regions which generate forebrain GABAergic neurons (Cobos et al., 2006; Flandin et al., 2010; Nobrega-Periera et al., 2010; Stanco et al., 2014). Many developmental transcription factors are also active in adulthood and are involved in maintaining the postmitotic identity of the neurons in which they are expressed (Deneris & Hobert, 2014). To determine the percentage of Npas1-tdTomato^+^ neurons which express NPAS1 protein in adults, we performed immunohistochemical staining for NPAS1 protein in adult Npas1-cre-2A-tdTomato mice (n=3). As in our other experiments using these mice, low levels of tdTomato were amplified using an anti-RFP stain. As expected for a transcription factor, staining for NPAS1 protein was observed in the nucleus but not the cytoplasm. Overall, the distribution of NPAS1 protein was similar to the distribution of tdTomato^+^ neurons and most tdTomato^+^ neurons in the BF of adult Npas1-cre-2A-tdTomato mice also expressed NPAS1 protein (75.49 ± 2.47%, **Fig. 1F**). Expression of NPAS1 was not significantly different across Npas1-tdTomato^+^ cells from rostral, medial, and caudal BF sections. Highest levels of colocalization were found in the HDB (87.07 ± 3.12%) and MCPO (80.53 ± 2.38%) with slightly lower levels found in the fiber-dense SI/VP (67.51 ± 5.13%), possibly due to poor antibody penetration in the region. Npas1-tdTomato^+^ neurons in regions neighboring the BF also expressed NPAS1 protein, as previously demonstrated in the globus pallidus pars externa (Hernandez et al., 2015).

### The vast majority of BF Npas1^+^ neurons are GABAergic but do not express parvalbumin, choline acetyltransferase or vesicular glutamate transporters

In other forebrain areas such as the cortex, hippocampus and GPe, Npas1 is expressed in GABAergic neurons (Stanco et al., 2014; Hernandez et al., 2015). To test if this is also the case in the BF, we performed immunohistochemistry for NPAS1 protein in neurons expressing fluorescent GABAergic (GAD67-GFP knock-in mice; Tamamaki et al., 2003; McKenna et al., 2013) and glutamatergic markers (vGlut1-cre-tdTomato and vGlut2-cre-tdTomato mice). In GAD67-GFP knock-in mice (n=3) the vast majority of NPAS1^+^ nuclei colocalized with GFP fluorescence (**Fig. 2A**). Overall: 98.24 ± 0.33% of NPAS1^+^ nuclei colocalized with GFP (SI/VP: 96.58 ± 0.43%; HDB: 96.84 ± 1.90%; MCPO: 99.05 ± 0.67%). No significant differences were found between rostral, medial and caudal sections. Similarly high colocalization was also observed in one GAD67-GFP/Npas1-cre-2A-tdTomato crossed mouse (**Supplemental Figure 1D**).

**Figure 2.**
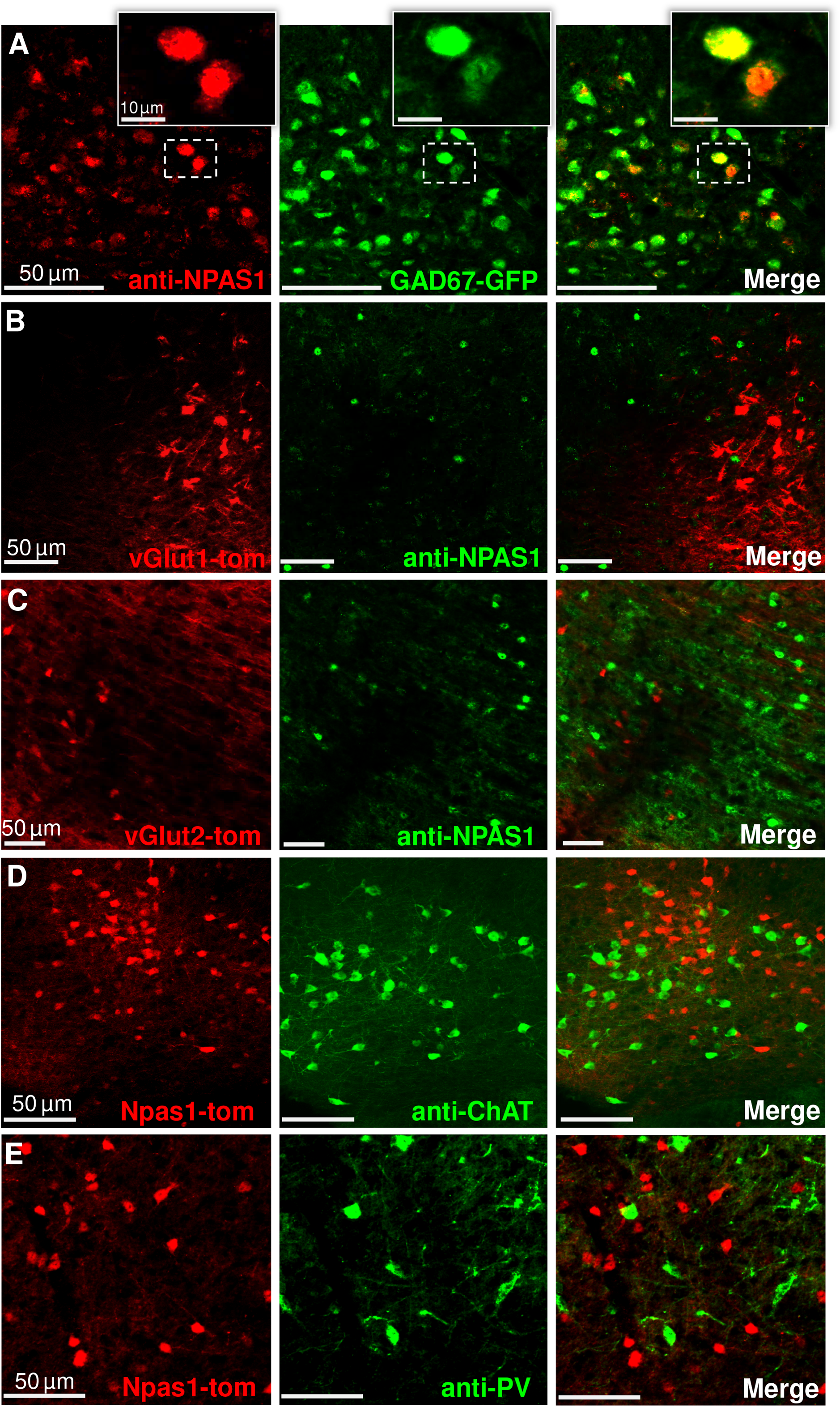
Npas1-expressing neurons of the BF are GABAergic neurons distinct from parvalbumin (PV) and cholinergic (choline acetyltransferase, ChAT) neurons. **(A)** The vast majority of IHC stained NPAS1^+^ nuclei (red) colocalize with green fluorescent protein (GFP) of GAD67-GFP knock-in mice. NPAS1^+^ nuclei (green) do not colocalize with tdTomato (red) in **(B)** vGlut1-cre-tdTomato or **(C)** vGlut2-cre-tdTomato mice. **(D, E)** Npas1-tdTomato+ neurons (red) do not colocalize with neurons stained against ChAT (green, **D**) or PV (green, **E**).

In vGlut1-cre-tdTomato (n=3) and vGlut2-cre-tdTomato mice (n=5), anti-NPAS1 immunohistochemistry revealed minimal (<0.5%) colocalization of NPAS1 protein (green) with red fluorescent glutamatergic neurons (**Fig. 2B, C**). To determine if Npas1^+^ neurons express the cholinergic marker choline acetyltransferase (ChAT) or the calcium-binding protein PV, we performed immunohistochemistry in Npas1-cre-2A-tdTomato mice (n=3 animals). Looking throughout the BF, there was similarly low colocalization between PV^+^ or ChAT^+^ neurons with Npas1-tdTomato^+^ neurons (**Fig. 2D, E**). As described above, Npas1-tdTomato^+^ neurons also tended to be smaller than most PV^+^ or cholinergic neurons. Thus, together, our immunohistochemical data suggest that BF Npas1^+^ neurons represent a subset of non-cholinergic, GABAergic neurons distinct from BF PV^+^ neurons.

### Anterograde tracing using AAV5-DIO-ChR2-EYFP injections revealed projections of BF Npas1^+^ neurons to brain areas involved in sleep-wake control, motivated behavior, and olfaction

To determine the targets of BF Npas1^+^ neurons, we performed anterograde tracing experiments using unilateral injections of AAV5-DIO-ChR2-EYFP into the BF of adult Npas1-cre-tdTomato mice to specifically express ChR2-EYFP in BF Npas1^+^ neurons and their efferent projections (**Fig. 3**). The EYFP fluorescence was amplified with an anti-GFP IHC stain to more easily identify transduced axons of BF Npas1^+^ neurons. Post-hoc analysis revealed good targeting of AAV5-DIO-ChR2-EYFP infusions to the ventral BF (HDB and MCPO) and dorsal SI/VP regions (n=6 mice; **Figs. 3A, B**). There was some limited transduction in a region medial to the BF which, in our previous studies and in the mouse brain atlas of Franklin and Paxinos, is labeled substantia innominata, but is included within the lateral preoptic area in the Allen Brain Atlas. Since this region is an area which contains cholinergic projection neurons (McKenna et al., 2013), it can be considered as part of the BF, as conventionally described.

**Figure 3.**
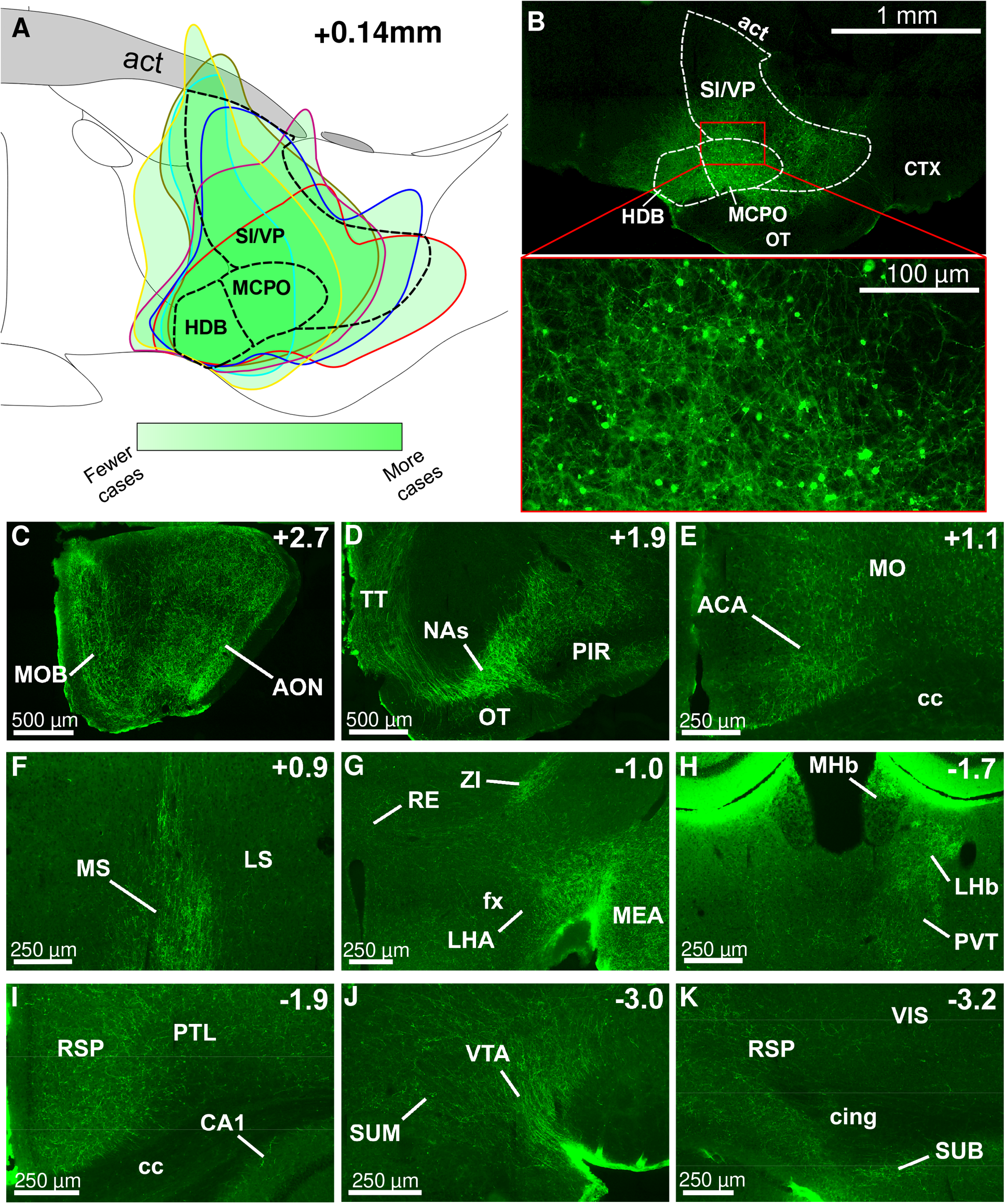
BF Npas1^+^ neurons project to brain areas involved in sleep-wake control, motivated behavior and olfaction. **(A)** Composite schematic showing the boundaries of unilateral AAV5-DIO-ChR2-EYFP infusions in the medial BF (from bregma, AP +0.14 mm; ML +1.6 mm; DV −5.3 mm) in 6 mice, with darker green indicating more frequent transduction of the region. Transduced neurons were largely restricted to the BF. **(B)** Low (top) and high (bottom) magnification images of an individual injection. Abbreviations: act, anterior commissure temporal limb; CTX, cortex; OT, olfactory tubercle. (**C-K)** Rostral to caudal images of regions with prominent fiber projections. Top right indicates position relative to bregma. **(C, D)** BF Npas1^+^ cells strongly project to olfactory regions, including main and anterior olfactory nuclei (MOB, AON), taenia tecta (TT) and piriform cortex (PIR). **(D)** Fibers largely avoid the core of the nucleus accumbens, but project to the shell (NAs). **(E, I, K)** Projections were dispersed throughout the neocortex, with denser projections in the anterior cingulate area (ACA), motor cortex (MO), retrosplenial cortex (RS), endopiriform nucleus and infralimbic and prelimbic cortices. **(F)** Fibers are widespread throughout septal regions rostral to the BF. Medial septal (MS) fibers were notably denser than those to the lateral septum (LS). **(G)** Hypothalamic projections were strongest in the lateral hypothalamic area (LHA) with moderate projections to the zona incerta (ZI), posterior, anterior, and dorsomedial hypothalamic nuclei and **(J)** supramammillary nucleus (SUM). **(G)** Dense projections were also observed in the amygdala, including the medial amygdalar nucleus (MEA), as well as the central amygdalar nucleus and cortical amygdala. **(H)** Projections were diffuse throughout the thalamus, highest in the lateral habenula (LHb) and paraventricular nucleus (PVT). **(H, I, K)** Fibers were present throughout most hippocampal formations, but strongest in the subiculum (SUB) and entorhinal cortex. **(J)** Densest midbrain projections were to the ventral tegmental area (VTA), with sparser projections to interpeduncular and pedunculopontine nuclei, the dorsal and medial raphe and the periaqueductal grey.

Anterograde tracing revealed prominent forebrain and midbrain projections of BF Npas1^+^ neurons (**Fig. 3**, **Table 1**). Projections were predominantly ipsilateral (**Table 1**). In the neocortex, labeled fibers were observable in almost all regions but were generally sparse, with the exception of moderate fiber densities in the frontal cortex (prelimbic, infralimbic, anterior cingulate and endopiriform cortices; **Fig. 3E**), and in retrosplenial cortex (**Figs. 3I, 3K**). Fibers were also evident in most subregions of the hippocampal formation (**Figs. 3I, K**), most prominently in the entorhinal cortex and dorsal subiculum (**Fig. 3K**). The strongest projections to the thalamus targeted the lateral habenula (**Fig. 3H**) and the paraventricular thalamic nucleus (**Fig. 3H**), a stress-related region which also regulates wakefulness (Ren et al., 2018), although scattered fibers were also present in other associative and sensory/motor relay nuclei (**Table 1**). There were only weak projections to the thalamic reticular nucleus, in marked contrast to the strong projections of GPe Npas1^+^ neurons (Abecassis et al., 2020; Cui et al., 2021), as well as BF PV^+^ and cholinergic neurons (Hallanger et al., 1987; Anaclet et al., 2015; Do et al., 2016; Thankachan et al., 2019). There were high fiber densities in rostral BF regions such as the medial septum, septofimbrial nucleus and triangular nucleus of the septum, and a moderate density of fibers in the vertical limb of the diagonal band, but only low levels in the lateral septum (**Fig. 3F**; **Table 1**). Amygdalar regions also tended to have high or moderate fiber densities (**Fig. 3G**, **Table 1**). A prominent finding was very high fiber densities in brain areas involved in olfaction, including the main olfactory bulb (**Fig. 3C**), anterior olfactory nucleus (**Fig. 3C**) and taenia tecta (**Fig. 3D**). In the basal ganglia, a strong projection was observed in the shell of the nucleus accumbens (**Fig. 3D**) as well as to the midbrain ventral tegmental area (**Fig. 3J**). BF Npas1^+^ fibers largely avoided the core of the nucleus accumbens (**Fig. 3D**) and the dorsal striatum (**Table 1**). In the hypothalamus, the strongest projection was to the lateral hypothalamic area (**Fig. 3G**) which contains orexin/hypocretin and other wake-promoting neuronal phenotypes (Arrigoni et al., 2019). The lateral supramammillary nucleus (**Fig. 3J**), posterior hypothalamus and zona incerta (**Fig. 3G**) exhibited a moderate density of fibers, as did the anterior hypothalamic nucleus, dorsomedial hypothalamus, and medial preoptic area (**Table 1**). Few fibers were observed in sleep-promoting areas such as the ventrolateral preoptic area and median preoptic area. The fiber density was generally low in the pons, but fibers could be observed in sleep-wake regulatory regions such as the raphe nuclei, pedunculopontine nucleus and periaqueductal gray (**Table 1**).

**Table 1.**
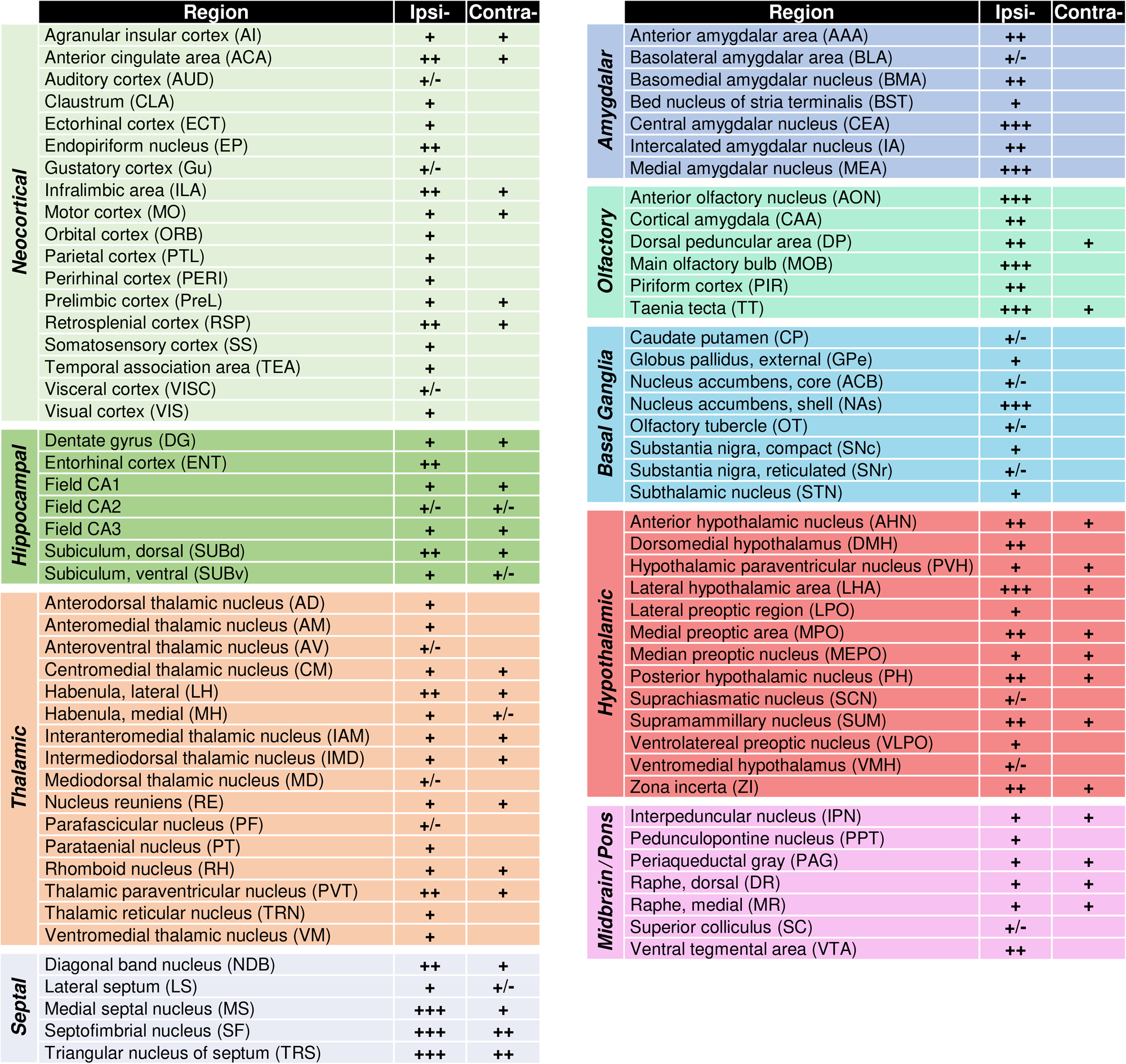
Comprehensive list of BF Npas1^+^ neuron ipsi- and contralateral projections within major brain regions and subregions. +/−, minimal or ambiguous fibers. +, ++, +++, low, moderate, and high density of terminal fibers, respectively. The density of contralateral projections is reported if observed.

|  | Region | Ipsi- | Contra- |
| --- | --- | --- | --- |
| Neocortical | Agranular insular cortex (AI) | + | + |
|  | Anterior cingulate area (ACA) | ++ | + |
|  | Auditory cortex (AUD) | +/- |  |
|  | Clastrum (CLA) | + |  |
|  | Ectorhinal cortex (ECT) | + |  |
|  | Endopiriform nucleus (EP) | ++ |  |
|  | Gustatory cortex (Gu) | +/- |  |
|  | Infralimbic area (ILA) | ++ | + |
|  | Motor cortex (MO) | + | + |
|  | Orbital cortex (ORB) | + |  |
|  | Parietal cortex (PTL) | + |  |
|  | Perirhinal cortex (PERI) | + |  |
|  | Prelimbic cortex (PreL) | + | + |
|  | Retrosplenial cortex (RSP) | ++ | + |
|  | Somatosensory cortex (SS) | + |  |
|  | Temporal association area (TEA) | + |  |
|  | Visceral cortex (VISC) | +/- |  |
|  | Visual cortex (VIS) | + |  |
| Hippocampal | Dentate gyrus (DG) | + | + |
|  | Entorhinal cortex (ENT) | ++ |  |
|  | Field CA1 | + | + |
|  | Field CA2 | +/- | +/- |
|  | Field CA3 | + | + |
|  | Subiculum, dorsal (SUBd) | ++ | + |
|  | Subiculum, ventral (SUBv) | + | +/- |
| Thalamic | Anterodorsal thalamic nucleus (AD) | + |  |
|  | Anteromedial thalamic nucleus (AM) | + |  |
|  | Anteroventral thalamic nucleus (AV) | +/- |  |
|  | Centromedial thalamic nucleus (CM) | + | + |
|  | Habenula, lateral (LH) | ++ | + |
|  | Habenula, medial (MH) | + | +/- |
|  | Interanteromedial thalamic nucleus (IAM) | + | + |
|  | Intermediodorsal thalamic nucleus (IMD) | + | + |
|  | Mediodorsal thalamic nucleus (MD) | +/- |  |
|  | Nucleus reuniens (RE) | + | + |
|  | Parafascicular nucleus (PF) | +/- |  |
|  | Parataenial nucleus (PT) | + |  |
|  | Rhomboid nucleus (RH) | + | + |
|  | Thalamic paraventricular nucleus (PVT) | ++ | + |
|  | Thalamic reticular nucleus (TRN) | + |  |
|  | Ventromedial thalamic nucleus (VM) | + |  |
| Septal | Diagonal band nucleus (NDB) | ++ | + |
|  | Lateral septum (LS) | + | +/- |
|  | Medial septal nucleus (MS) | +++ | + |
|  | Septofimbrial nucleus (SF) | +++ | ++ |
|  | Triangular nucleus of septum (TRS) | +++ | ++ |
| Amygdalar | Anterior amygdalar area (AAA) | ++ |  |
|  | Basolateral amygdalar area (BLA) | +/- |  |
|  | Basomedial amygdalar nucleus (BMA) | ++ |  |
|  | Bed nucleus of stria terminalis (BST) | + |  |
|  | Central amygdalar nucleus (CEA) | +++ |  |
|  | Intercalated amygdalar nucleus (IA) | ++ |  |
|  | Medial amygdalar nucleus (MEA) | +++ |  |
| Olfactory | Anterior olfactory nucleus (AON) | +++ |  |
|  | Cortical amygdala (CAA) | ++ |  |
|  | Dorsal peduncular area (DP) | ++ | + |
|  | Main olfactory bulb (MOB) | +++ |  |
|  | Piriform cortex (PIR) | ++ |  |
|  | Taenia tecta (TT) | +++ | + |
| Basal Ganglia | Caudate putamen (CP) | +/- |  |
|  | Globus pallidus, external (GPe) | + |  |
|  | Nucleus accumbens, core (ACB) | +/- |  |
|  | Nucleus accumbens, shell (NAs) | +++ |  |
|  | Olfactory tubercle (OT) | +/- |  |
|  | Substantia nigra, compact (SNc) | + |  |
|  | Substantia nigra, reticulated (SNr) | +/- |  |
|  | Subthalamic nucleus (STN) | + |  |
| Hypothalamic | Anterior hypothalamic nucleus (AHN) | ++ | + |
|  | Dorsomedial hypothalamus (DMH) | ++ |  |
|  | Hypothalamic paraventricular nucleus (PVH) | + | + |
|  | Lateral hypothalamic area (LHA) | +++ | + |
|  | Lateral preoptic region (LPO) | + |  |
|  | Medial preoptic area (MPO) | ++ | + |
|  | Median preoptic nucleus (MEPO) | + | + |
|  | Posterior hypothalamic nucleus (PH) | ++ | + |
|  | Suprachiasmatic nucleus (SCN) | +/- |  |
|  | Supramammillary nucleus (SUM) | ++ | + |
|  | Ventrolatereal preoptic nucleus (VLPO) | + |  |
|  | Ventromedial hypothalamus (VMH) | +/- |  |
|  | Zona incerta (ZI) | ++ | + |
| Midbrain / Pons | Interpeduncular nucleus (IPN) | + | + |
|  | Pedunculo pontine nucleus (PPT) | + |  |
|  | Periaqueductal gray (PAG) | + | + |
|  | Raphe, dorsal (DR) | + | + |
|  | Raphe, medial (MR) | + | + |
|  | Superior colliculus (SC) | +/- |  |
|  | Ventral tegmental area (VTA) | ++ |  |

In summary, these cell-specific anterograde experiments revealed that BF Npas1^+^ cells are projection neurons that strongly innervate subcortical regions and have a dual role in control of sleep-wake regulation and motivated behavior. Additionally, there are strong projections of BF Npas1^+^ neurons to brain areas involved in olfaction. Given previous studies of BF GABAergic neurons and the projections of BF Npas1^+^ neurons to sleep-wake control regions, we next determined the effect of activating BF Npas1^+^ neurons on sleep-wake behavior and cortical oscillations.

### Chemogenetic stimulation of BF Npas1^+^ neurons promotes wakefulness and disrupts NREM slow and sigma oscillations

To determine the functional effect of activating BF Npas1^+^ neurons on sleep-wake behavior and cortical EEG, we performed chemogenetic experiments in Npas1-cre-2A-tdTomato mice. Viral vectors for Cre-dependent expression of the excitatory receptor hM3d(Gq) and a fluorescent marker of transduction (mCherry) were injected bilaterally into the BF (**Fig. 4A, B**). 4 or more weeks later, mice received intraperitoneal (i.p.) injections of saline, 0.3 mg/kg or 1 mg/kg CNO in a within-subject design to activate BF Npas1^+^ neurons at the beginning of the light (inactive) period at zeitgeber time 2 (ZT2). Post-hoc histological examination confirmed successful targeting of the BF (**Fig. 4A, B**) and transduction of Npas1^+^ neurons (**Fig. 4C**). Limited transduction was observed in preoptic areas medial to the BF. In a subset of cases (2/9 cases bilaterally and 2/9 unilaterally), there was spread dorsal to the anterior commissure. However, results were comparable to those cases without spillover across the anterior commissure and have been combined for the analysis. Following saline injections (n=9 mice), average sleep latency was <30 min and REM latency was <50 min. In contrast, injections of 0.3 mg/kg (n=9) or 1 mg/kg CNO (n=7) resulted in significantly longer latencies to both non-REM sleep (approximately 100 min) and REM sleep (>150 min) (**Fig. 4D**). Overall, compared to saline (27.2 ± 5.7 min), NREM sleep latency was significantly higher following administration of 0.3 mg/kg (98.8 ± 17.5 min; p=0.0045) or 1 mg/kg CNO (105.0 ± 21.5 min; p=0.0045, one-way RM ANOVA, **Fig. 4E**). Similarly, in comparison to saline administration (47.0 ± 5.7 min), latency to REM sleep was also significantly higher following injections of 0.3 mg/kg CNO (155.3 ± 14.2 min; p<0.0001) and 1 mg/kg CNO (181.6 ± 23.7 min; p<0.0001, one-way RM ANOVA, **Fig. 4F**).

**Figure 4.**
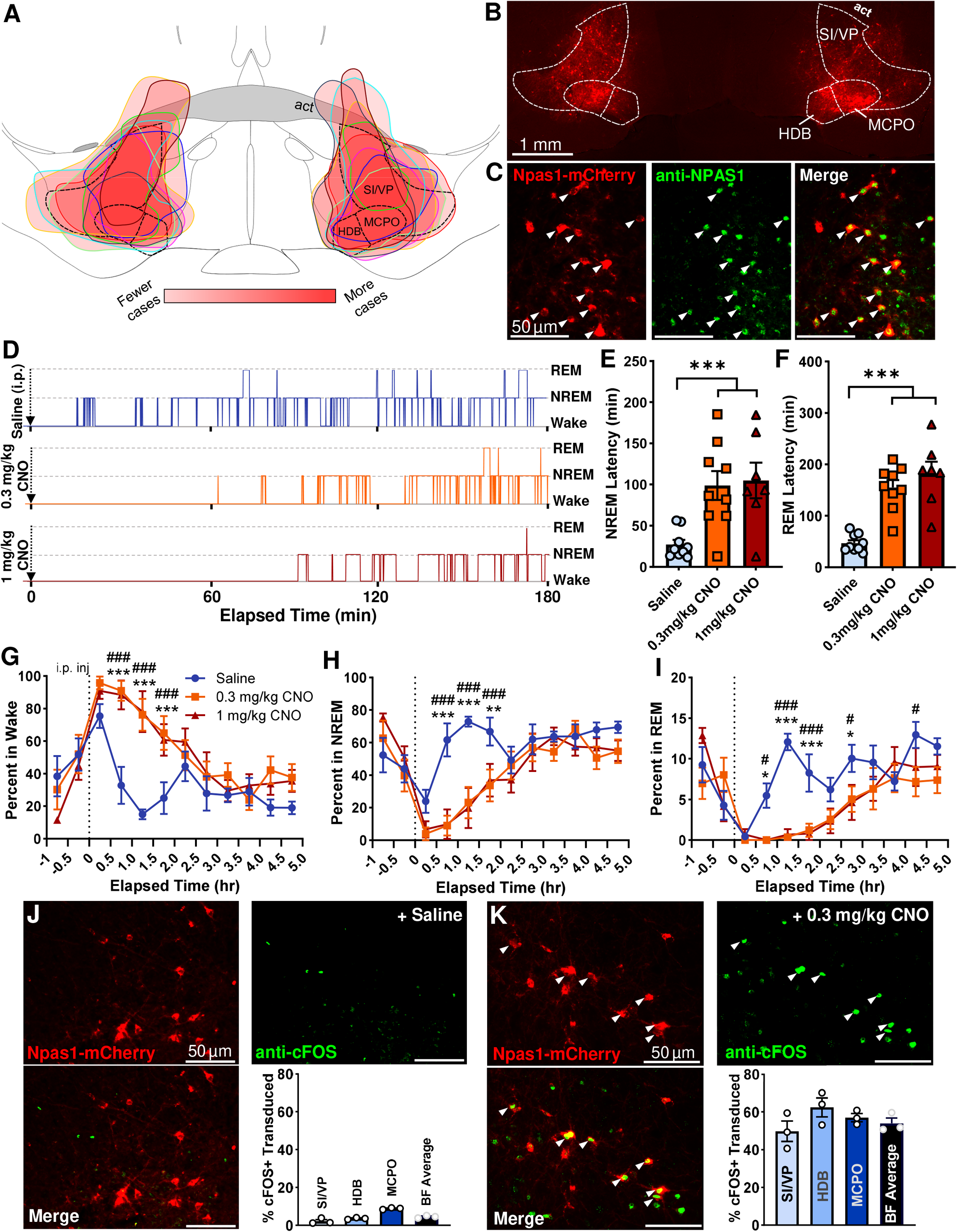
Activation of BF Npas1^+^ neurons promotes wakefulness and suppresses NREM and REM sleep. **(A)** Injection composite indicating the spread of individual bilateral pAAV-hSyn-DIO-hM3D(Gq)-mCherry infusions in the medial BF (from bregma, AP +0.14 mm; ML +1.6 mm; DV −5.3 mm) across 9 animals. Darker red indicates more frequent targeting of the region. **(B)** Example bilateral infusion of pAAV-hSyn-DIO-hM3D(Gq)-mCherry in a medial BF section. **(C)** The majority of transduced Npas1-cre cells expressing mCherry (red) also colocalize with an NPAS1 nuclear stain (green), indicating most transduced Npas1-cre-tdTomato^+^ neurons had active Npas1 expression in adults. **(D)** Representative hypnogram from one mouse following i.p. injections of 0.3 mg/kg CNO (orange), 1 mg/kg CNO (red) or saline (blue). The onset of NREM sleep is <30 min following saline administration, but >60 min after CNO administration. Latency to REM sleep following saline injection is <90 min, but >150 min after CNO. **(E)** Compared to saline (n=9), NREM latency was significantly higher following administration of 0.3 mg/kg CNO (n=9) or 1 mg/kg CNO (n=7). A one-way ANOVA finds a significant effect of treatment on NREM latency (F_2,22_ = 8.02 p=0.0024). **(F)** Additionally, REM latency was significantly longer following injections of 0.3 mg/kg CNO and 1 mg/kg CNO compared to saline. One-way ANOVA finds significant effect of treatment on REM latency (F_2,22_ = 23.2 p<0.0001). **(G)** Administration of 0.3 or 1 mg/kg CNO promotes significantly more wakefulness vs. saline beginning 0.5h after injection and persisting for approximately 1.5h. Two-way ANOVA identifies significant interaction of time x treatment (F_22,264_ = 2.69, p=0.0001). **(H)** Concomitantly, 0.3 and 1 mg/kg CNO injections promote significantly less time in NREM sleep vs. saline. Two-way ANOVA identifies significant interaction of time x treatment (F_22,264_ = 2.60, p=0.0002). **(I)** REM sleep is also significantly suppressed after CNO administration vs. saline, starting 0.5h post-injection and persisting approximately 3 hours. Two-way ANOVA identifies significant interaction of time x treatment (F_22,264_ = 2.66, p=0.0001). Saline vs. 0.3 mg/kg CNO, p <0.05 #, p<0.01 ##, p<0.005 ###. Saline vs 1 mg/kg CNO: p<0.05 *, p<0.01 **, p<0.005 ***. **(J)** Few BF NPAS1-mCherry cells (red) expressed cFOS following saline administration 2h prior to sacrifice (4.5% ± 0.4%, n=3). **(K)** Colocalization with immunohistochemical nuclear stains for cFOS (green) indicate the majority of transfected BF NPAS1-mCherry cells (red) were activated by 0.3 mg/kg CNO administered 2h prior to sacrifice (54.0% ± 2.8%, n=3), with no difference between subregions. Colocalization is marked with an arrowhead. p<0.005 ***, ns: not significant.

I.P. administration of 0.3 mg/kg and 1 mg/kg CNO significantly increased the amount of wakefulness and suppressed NREM and REM sleep vs. saline in the first 2-3 hours post-injection (**Fig. 4G, H, I**). Two-way RM ANOVA indicated a significant interaction of time and treatment on the proportion of time spent in wake (F_22, 264_ =2.69, p=0.0001), NREM (F_22, 264_ = 2.60, p=0.0002) and REM (F_22, 264_ = 2.66, p=0.0001). Latencies and amounts of wakefulness, NREM or REM sleep were not significantly different between 0.3 and 1mg/kg doses of CNO. 0.3 mg/kg CNO alone had no significant effect on sleep-wake behavior or cortical EEG in a subset of mice of the same background strain where Npas1^+^ neurons were not transduced with hM3d(Gq) (n=4, **Supplemental Figure 2**).

A subset of mice was given saline, 0.3 or 1 mg/kg CNO at ZT2, 2 hours prior to sacrifice to confirm activation of BF Npas1-mCherry^+^ neurons by CNO using IHC for the immediate early gene product cFOS. In saline treated mice (n=3; **Fig. 4K**), the percentage of mCherry^+^ neurons expressing cFOS was <5% (4.5 ± 0.4%, n=3), whereas following 0.3 mg/kg CNO (**Fig. 4J**) it was >50% (54.0 ± 2.8%, n=3; p<0.0001, two-tailed unpaired t-test), confirming strong activation of the Npas1^+^ neurons by CNO (**Fig 4J, K**). Similarly high percentages were seen following 1 mg/kg CNO (57.2 ± 3.9%; n=2). The percentage of neighboring PV^+^ and cholinergic (ChAT^+^) neurons which expressed cFOS also increased following CNO injections (**Supplemental Figure 3**), consistent with the wake-active profile of these neurons (Anaclet et al., 2015; Xu et al., 2015; McKenna et al., 2020), but remained below 20% (PV; 4.7 ± 1.5% with saline, n=3; 16.4 ± 3.7% with 0.3 mg/kg CNO, n=3; p<0.05, two-tailed unpaired t-test) and 10%, respectively (ChAT; 2.2 ± 1.1% with saline, n=3; 5.7 ± 0.6% with 0.3 mg/kg CNO, n=3; p=0.05, two-tailed unpaired t-test).

To further explore the increase in wakefulness following chemogenetic activation of BF Npas1^+^ neurons, we performed sleep-wake bout analysis. One mouse treated with 0.3 mg/kg and its saline control data was excluded since it exhibited continuous wakefulness for >3 hr post-injection of CNO and was identified as an outlier by a Grubbs’ outlier test (p<0.0001). In the first 3 hrs post-injection, the duration of wake bouts was increased by 0.3 mg/kg CNO (Saline 58.4 ± 5.4 s; 0.3 mg/kg 104.3 ± 18.0 s; p=0.021, two-tailed paired t-test, **Fig 5A**) and by 1 mg/kg CNO (213.2 ± 70.5 s, n=7; p=0.0039, two-tailed paired t-test, **Supplemental Figure 4A**). NREM bout duration was reduced by 0.3 mg/kg CNO (saline 95.9 ± 5.9 s; 0.3 mg/kg CNO 40.3 ± 9.3 s; p=0.002, two-tailed paired t-test, **Fig. 5A**) and tended to be reduced by 1 mg/kg CNO (58.5 ± 22.2 s; p=0.14, two-tailed paired t-test, **Supplemental Figure 4A**). The number of wake and NREM bouts was unaltered (**Fig. 5B**). REM bout duration was not significantly altered by 0.3 mg/kg (**Fig. 5A**) or 1 mg/kg CNO (**Supplemental Figure 4A**), but the number of REM bouts was strongly and significantly reduced (p<0.001, two-tailed paired t-test, **Fig. 5B**, **Supplemental Figure 4B**). Neither dose of CNO significantly changed the number of brief awakenings, defined as four or fewer epochs (<16 s) of wakefulness interrupting 2 sleep bouts (**Fig. 5C, Supplemental Figure 4C**). Further analysis of individual bouts during the 3 hr post-injection period revealed that following 0.3 mg/kg CNO administration, shorter wake bouts were less frequent (**Fig. 5D**), with significantly fewer wakefulness bouts between 13-20 s (p=0.015) and significantly more exceeding 400 s (p=0.020, two-tailed paired t-test, **Fig. 5G**). Shorter NREM bouts are more frequent following 0.3 mg/kg CNO administration (**Fig 5E**), producing significantly more NREM bouts shorter than 20 s (p<0.05) and reducing those longer than 100 s (p<0.05, two-tailed paired t-test, **Fig. 5H**). Across treatments, the distribution of REM bout lengths was not significantly different (**Fig 5F**), although 0.3 mg/kg CNO significantly decreased REM bouts between 13-100 s in length (p<0.05, two-tailed paired t-test, **Fig. 5I**). Trends in cumulative wake, NREM, and REM bout length distributions and bout frequency comparisons were comparable following administration of 1 mg/kg CNO (**Supplementary Figure 4 D-I**). Overall, these data suggest that increases in the percentage of time in wakefulness and corresponding decreases in NREM time caused by chemogenetic activation of BF Npas1^+^ neurons are due primarily to an increase in long-duration bouts of wakefulness and an increase in short duration NREM bouts. In contrast, suppression of percentage time in REM sleep is due to a suppression of entries into REM.

**Figure 5.**
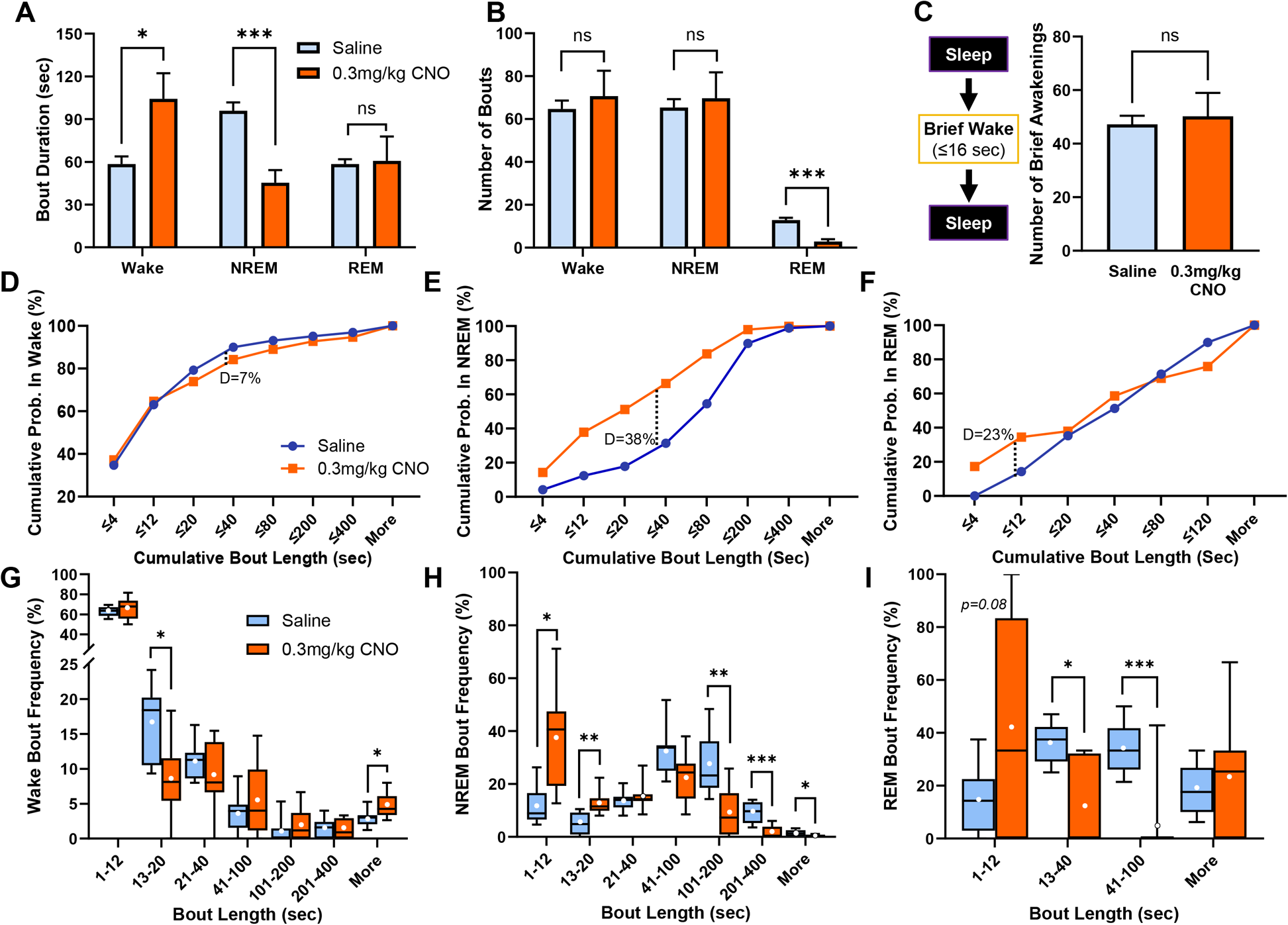
Activation of BF Npas1^+^ cells increases the duration of wake bouts while reducing the duration of NREM bouts and number of REM bouts. **(A)** Compared to saline (n=8), administration of 0.3 mg/kg CNO (n=8) produced significantly longer wake bouts (p=0.021) and significantly shorter NREM bouts (p=0.0020) over the 3-hour period post-injection period. REM bout length was unaffected (p=0.89). Comparisons were made via two-tailed paired t-test **(B)** Administration of 0.3 mg/kg CNO did not change the number of wake or NREM bouts. However, there were significantly fewer REM bouts after 0.3 mg/kg CNO administration (p=0.00013). p<0.05*, p<0.005 ***, ns: not significant). **(C)** CNO administration did not change the number of brief awakenings, defined as ≤4 epochs (≤16 sec) of wakefulness interspersing epochs of NREM sleep (p=0.71). Cumulative frequency plots of bouts **(D, E, F)** and histograms of bout length **(G, H, I)** during wake, NREM and REM respectively identify further effects on sleep architecture. **(D)** Cumulative probability of wake during the 3-hour period following 0.3 mg/kg CNO administration trends toward less frequent short wake bouts (p=0.093, K-S D=0.070). **(G)** Bout analysis indicates significantly fewer wake bouts between 13-20 seconds long (p=0.015) and more frequent bouts over 400 seconds (p=0.020). **(E)** Shorter NREM bouts are more frequent following 0.3 mg/kg CNO administration, producing a significantly different cumulative frequency distribution (p<0.001, K-S D=0.38). **(H)** Bout analysis indicates CNO produces significantly more bouts between 1-12 and 13-20 seconds (p=0.015 and p=0.0098, respectively), and significantly fewer longer bouts between 101-200, 201-400, and 400+ seconds (p=0.0078, p=0.0006 and 0.018, respectively). **(F)** Overall cumulative distributions of REM bouts were not significantly different across treatments (p=0.16, K-S D=0.23), **(I)** Administration of 0.3 mg/kg CNO produced significantly fewer moderate-length REM bouts between 13-40 and 41-100 seconds (p=0.016 and p=0.0007, respectively). Cumulative histograms are compared as two-sample nonparametric Kolmogorov-Smirnov (K-S) tests, and individual binned bout comparisons were made via two-tailed paired t-tests. Whiskers on box and whisker plots range from min to max, the midline represents the median, and the mean is indicated with a white circle. Up to 1 animal per group bin may be excluded if it is alone in spending 0% or 100% of its time in that bin. p<0.05 *, p<0.01 **, p<0.005 ***, ns: not significant.

Next, we analyzed whether activation of BF Npas1^+^ neurons changed the quality of sleep-wake states, in terms of their spectral profile (**Fig. 6, Supplemental Figure 5**). During wakefulness, 0.3 mg/kg CNO in mice expressing hM3d(Gq) in BF Npas1^+^ neurons produced a significant increase in power in the theta (4-9 Hz, 115% of saline control, p=0.0043), alpha (9-15 Hz bands, to 121% of control, p=0.0051) and beta bands (15-30 Hz, 109% of control, p=0.039; two-tailed paired t-tests, **Fig. 6C**). Maximal differences compared to saline were observed at 6.8-7.3 Hz in the theta band (127% of control), 11.2-11.7 Hz in the alpha band (126% of control) and 25.4-25.9 Hz in the beta band (114% of control). Effects were similar but not as pronounced with 1 mg/kg CNO (115% of control in theta power, p=0.22; 114% of control in alpha power, p=0.090; 104% of control in beta power, p=0.14, two-tailed paired t-test, **Supplemental Figure 5C**). In NREM sleep, 0.3 mg/kg CNO led to significant decreases in slow-wave activity (0-1.5 Hz, to 60% of control values, p<0.001) and power in the sigma band (9-15 Hz, to 76% of control values; p=0.0098, two-tailed paired t-test, **Fig. 6D**). Maximal spectral differences during NREM were seen between 0.49-0.98 Hz (56% of control) in the slow-wave band and 11.2-11.7 Hz (70% of control in the sigma band). Effects on slow-wave and sigma power were also observed following 1 mg/kg CNO (52.9% of control in the slow band, p=0.004; 75.9% of control for the sigma band; p=0.0136, two-tailed paired t-test, **Supplemental Figure 5D**). Power spectral changes during REM sleep were less pronounced (**Fig. 6E**) with a smaller increase in the theta and gamma band power (30-80 Hz) with 0.3 mg/kg CNO (p<0.05, two-tailed paired t-test) which was not replicated with 1 mg/kg CNO (**Supplemental Fig. 5E**). Further analysis of sleep and NREM delta (0.5-4 Hz) power in the remainder of the light period did not identify any homeostatic sleep rebound in either measure (**Supplemental Fig. 5A, B**).

**Figure 6.**
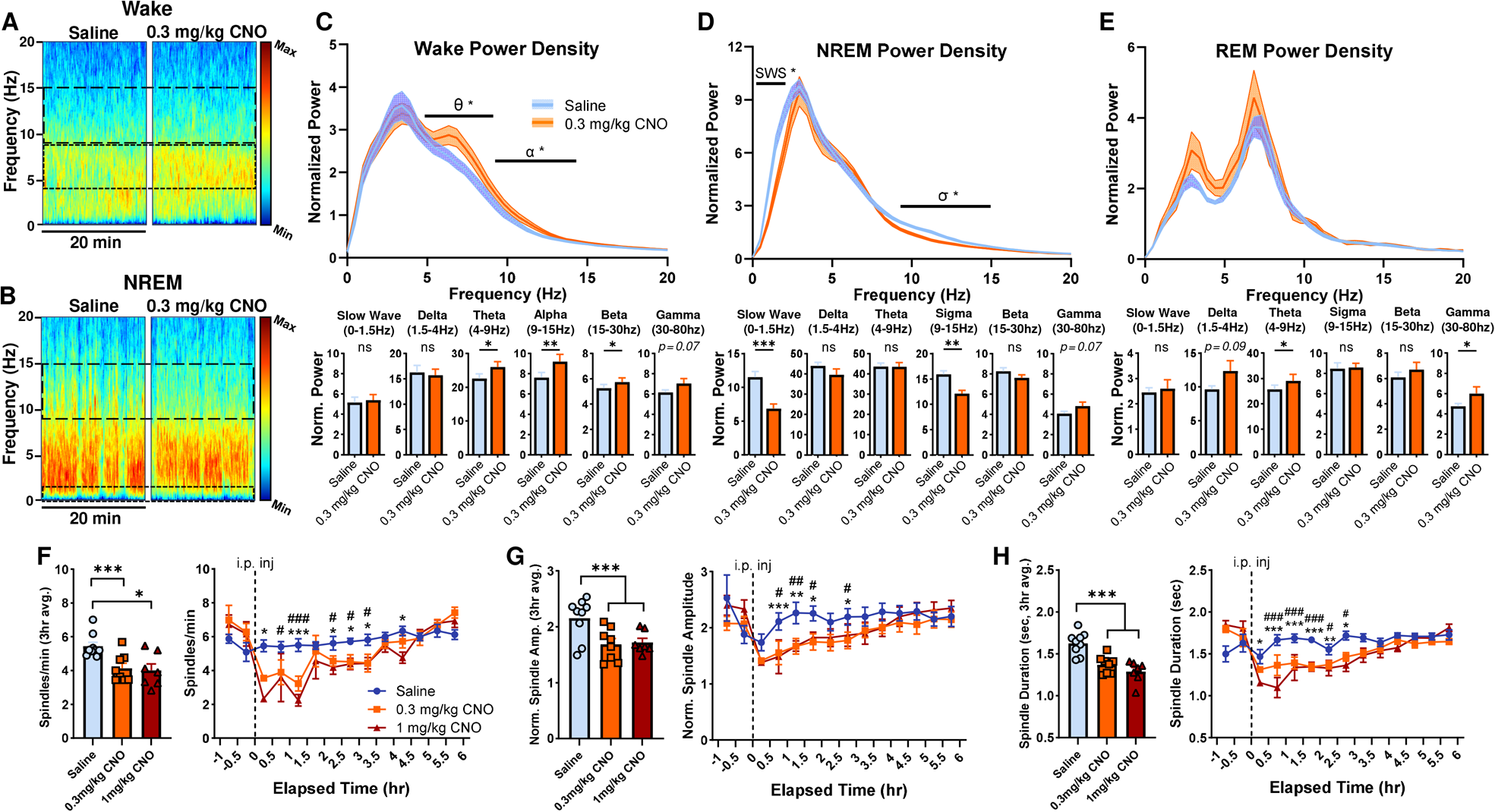
Chemogenetic activation of BF Npas1^+^ neurons alters cortical spectral components of sleep-wake states and suppresses NREM sleep spindles. **(A)** Representative EEG spectrogram during a continuous 20-minute sample of predominantly wakefulness occurring during the first hour following i.p. injection. More activity at a given frequency range is indicated by warmer colors, and less by cooler colors. Wake theta (4-9Hz) and alpha (9-15Hz) are outlined (dashed horizontal lines), and more relative activity in both bands was observed following 0.3 mg/kg CNO administration. **(B)** A continuous 20-minute excerpt of predominantly NREM EEG activity from the second post-treatment hour shows a decrease in NREM slow wave (0-1.5Hz) and sigma (9-15Hz) activity. **(C)** A normalized power spectral density plot between 0 and 20Hz during wakefulness (n=9 animals) indicates 0.3 mg/kg CNO produces a significant increase within theta and alpha bands during the 3-hour post-injection period, maximized at 6.8-7.3Hz and 11.2-11.7Hz, respectively. Individual power spectra normalized to total power and time in sleep-wake state finds significantly more activity in theta (p=0.0043), alpha (p=0.0051), and beta bands (p=0.039) but not slow wave (p=0.65), delta (p=0.58), or gamma (p=0.068). **(D)** Power spectra density during NREM indicates less power within the slow wave (0-1.5Hz) and sigma (10-15Hz) bands following 0.3 mg/kg CNO administration, with the greatest differences lying between 0.49-0.98Hz and 11.2-11.7Hz, respectively. Normalized power analysis concurrently finds significantly less slow wave (p<0.001) and sigma activity (p=0.0098), but no changes in delta (p=0.022), theta (p=0.82), beta (p=0.31) or gamma bands (p=0.067). **(E)** REM power spectra densities between 0-20Hz were not significantly different between saline or 0.3 mg/kg CNO groups, with no difference observed in slow wave (p=0.67), delta (p=0.092), sigma (p=0.60) or beta bands (p=0.25), but with a significant increase observed in the theta and gamma bands (p=0.034 and p=0.029, respectively). **(F)** Spindle density (NREM spindles/min) was significantly lower in mice administered 0.3 mg/kg (p=0.0004) or 1 mg/kg CNO (p=0.0321) vs. saline. Compared to saline, CNO produced significantly lower spindle densities within the first 0.5h after injection and persisting up to 3.5h. Mixed-effects analysis identifies significant interaction of time x treatment (F_26,139_ = 3.68, p=0.0001). **(G)** Normalized spindle amplitudes were significantly lower during the 3-hour post-injection period in mice administered 0.3 mg/kg (p=0.0048) or 1 mg/kg CNO (p=0.0048) vs. saline. Compared to saline, CNO produced significantly shorter spindle amplitudes beginning 1h after injection and persisting up to 3h. Mixed-effects analysis identifies interaction of time x treatment (F_26,139_ = 1.67, p=0.031). **(H)** Spindles were significantly shorter following administration of 0.3 mg/kg (p=0.0029) or 1 mg/kg CNO (p=0.0005) vs. saline. Spindles were significantly shorter within the first 0.5h post-injection, persisting up to 3h. Mixed-effects analysis identifies significant interaction of time x treatment (F_26,134_ = 4.58, p<0.0001). One animal administered 0.3 mg/kg CNO was excluded from 3-hour spindle analyses due to spending no time in NREM in this period. If an animal spent no time in NREM during an individual 30 min epoch in spindle time-course plots, that value of 0 was excluded from analysis. p<0.05 *, p<0.01 **, p<0.005 ***, ns: not significant.

Changes in NREM sigma power are often, but not always, indicative of changes in sleep spindles (Uygun et al., 2019; Astori et al., 2011), a feature of light NREM sleep associated with sleep stability and memory consolidation which is often impaired in many neuropsychiatric disorders (Fernandez & Luthi, 2020). Thus, we examined changes in sleep spindles after i.p. injections of saline or CNO using a recently validated algorithm (Uygun et al., 2019). Sleep spindle density, duration and amplitude were stable following saline injections, but large reductions in all three measures were observed during the 3-hour period following administration of 0.3 mg/kg or 1 mg/kg CNO (**Fig. 6F, G, H**). Spindle density (NREM spindles/min) was significantly lower in mice administered 0.3mg/kg CNO (4.14 ± 0.29 spindles/min; p=0.0004) or 1 mg/kg CNO (4.01 ± 0.38 spindles/min; p=0.0321, one-way RM ANOVA) vs. saline (5.44 ± 0.23 spindles/min) (**Fig. 6F**). Additionally, normalized spindle amplitudes were significantly smaller in mice administered 0.3 mg/kg (1.69 ± 0.10, p=0.0048) or 1 mg/kg CNO (1.72 ± 0.07; p=0.0048, one-way RM ANOVA) vs. saline (2.15 ± 0.12, **Fig. 6G**), as were spindle durations (**Fig. 6H**) after administration of 0.3 mg/kg (1.37 ± 0.037 s, p=0.0029) or 1 mg/kg CNO (1.29 ± 0.049 s; p=0.0005, one-way RM ANOVA) vs. saline (1.62 ± 0.045 s). These reductions were maximal within 60 min post-injection, lasted 3 or more hrs, and did not significantly differ between doses of CNO. No effects of 0.3 mg/kg CNO were found on spectral measures or sleep spindles in control mice without expression of hM3d(Gq) (n=4, **Supplemental Figure 6**).

In summary, chemogenetic activation of BF Npas1^+^ neurons strongly increased wakefulness, delayed the onset of sleep and disrupted NREM sleep oscillations, features reminiscent of sleep disruption in a variety of psychiatric and neurodegenerative disorders (Brown et al., 2022; Fernandez & Luthi, 2020).

## DISCUSSION

Here, we characterized a major subpopulation of BF projection neurons which express Npas1, a developmental transcription factor linked to neuropsychiatric disorders (Erbel-Sieler et al., 2004; Michaelson et al., 2017). Our results suggest that BF Npas1^+^ neurons are GABAergic, distinct from PV^+^ and cholinergic neurons, and project to brain regions involved in sleep-wake control, motivated behavior, and olfaction. When activated chemogenetically, BF Npas1^+^ neurons strongly enhance wakefulness, suppress NREM and REM sleep and disrupt NREM sleep oscillations, reminiscent of sleep disruption in a variety of disorders.

Although earlier studies reported Npas1 expression in the BF (Erbel-Sieler et al., 2004; Hernandez et al., 2015), ours is the first to comprehensively examine the location, density, size, and neurochemical phenotype of Npas1^+^ neurons in the three main BF subregions. In addition to Npas1 neurons in the VP (Morais-Silva et al., 2023), substantial numbers of small or medium-sized Npas1 neurons were located in the HDB and MCPO regions, as well as in neighboring lateral and medial preoptic areas. The density of Npas1^+^ neurons was substantially (5-6X) higher than of previously characterized BF PV^+^, ChAT^+^, and vGlut2^+^ neurons (McKenna et al., 2013, 2021; Yang et al., 2014). Immunohistochemical staining for NPAS1 revealed that the vast majority (∼75%) of BF Npas1-tdTomato^+^ neurons continue to express detectable levels of Npas1 into adulthood, likely an underestimate due to the fiber-dense nature of the BF which hinders antibody penetration. These results suggest a possible function for Npas1 in regulating the activity or phenotype of these neurons in adults, similar to other developmental transcription factors (Deneris & Hobert, 2014). IHC staining in Npas1-cre-2A-tdTomato^+^ tissue against PV and ChAT found negligible overlap, comparable to findings in the GPe and cerebral cortex (Nobrega-Pereira et al., 2010; Hernandez et al., 2015; Stanco et al., 2014) and consistent with the expression of Npas1 in embryonic precursors which generate forebrain GABAergic neurons (Erbel-Sieler et al., 2004; Cobos et al., 2006; Flandin et al., 2010; Nobrega-Pereira et al., 2010). In the BF, NPAS1^+^ nuclei colocalized with GFP in GAD67-GFP knock-in mice, whereas virtually no overlap was found between NPAS1 and tdTomato in vGlut1-cre-tdTomato or vGlut2-cre-tdTomato mice, indicating that Npas1^+^ neurons are not glutamatergic. Surprisingly, Morais-Silva et al., (2023) found that GABAergic markers were not enriched in VP Npas1^+^ neurons in a study which analyzed ribosome-associated mRNA in Npas1-cre-Ribotag mice, despite high expression of another transcription factor, Sox6, which is expressed in developing and adult forebrain GABAergic neurons (Azim et al., 2009; Abecassis et al., 2021). The origin of this discrepancy is unclear, but could reflect insufficient sensitivity of the Ribotag method. In contrast, our immunohistochemical staining experiments strongly suggest that BF Npas1^+^ neurons are GABAergic, like their counterparts in GPe and cortex.

To determine the projections of BF Npas1^+^ neurons, we expressed ChR2-EYFP fusion proteins in Npas1^+^ neurons and examined the location and relative density of green (EYFP^+^) fibers. In general, the results of this experiment were similar to those of a study (Morais-Silva et al., 2023) which focused on Npas1^+^ neurons in the VP, using the same Npas1-cre-2A-tdTomato mice. Both our study and that of Morais-Silva et al., (2023) revealed major projections to lateral hypothalamus, medial septum, nucleus accumbens, lateral habenula, VTA and anterior olfactory nucleus. However, our injections also transduced Npas1^+^ neurons in the HDB and MCPO BF subregions. We further identified additional projections to the neocortex, hippocampal formation, amygdala, supramamillary and posterior hypothalamus, and to the olfactory bulb. Overall, these results are consistent with BF Npas1^+^ neurons being involved in limbic basal ganglia circuits related to motivation, reward, stress and sleep-wake control (Luo et al., 2018; Li et al., 2021; Fifel et al., 2022; Morais-Silva et al., 2023; Soares-Cunha & Heinsbroek, 2023).

There are parallels between the previously described projections of Npas1^+^ neurons in the GPe involved in motor control and BF Npas1^+^ neurons involved in motivational circuitry. GPe Npas1^+^ neurons project to the dorsal striatum (Glajch et al., 2016), whereas BF Npas1^+^ neurons project strongly to ventral striatum (shell of the nucleus accumbens). GPe Npas1^+^ neurons project to the substantia nigra, whereas BF Npas1^+^ neurons project heavily to VTA. We also observed projections to the neocortex, especially frontal regions, similar to previously described neocortical projections of GPe Npas1^+^ neurons (Abecassis et al., 2020). One notable difference between the projections of GPe Npas1^+^ neurons and BF Npas1^+^ neurons was that we did not observe a strong projection of BF Npas1^+^ neurons to the TRN, whereas there is a strong output from GPe Npas1^+^ neurons (Abecassis et al., 2020; Cui et al., 2021). This finding also distinguishes BF Npas1^+^ neurons from BF PV^+^ and ChAT^+^ neurons which project heavily to the TRN (Hallanger et al., 1987; Do et al., 2016; Kim et al., 2019; Thankachan et al., 2019;).

The prominence of BF Npas1^+^ projections to olfactory regions was a surprising finding of this study. A major projection from the BF to the olfactory bulb (OB) and olfactory cortices has been known for some time and comprises both cholinergic and GABAergic components (Zaborszky et al., 1986; Sanz-Diez et al., 2019; Villar et al., 2021; Venegas et al., 2023). The cholinergic BF projection to olfactory bulb originates in the medial part of the HDB whereas the GABAergic component originates in lateral HDB and MCPO regions (Zaborszky et al., 1986; Gracia-Llanes et al., 2010). However, the identity of BF GABAergic neurons mediating this projection was unknown. Previous tracing studies found that BF PV+ and SST+ projections to OB are weak (Do et al., 2016). In contrast, our results suggest that BF GABAergic neurons of the Npas1^+^ lineage are a major component of this projection, with strong projections to not only OB but also olfactory cortex regions such as the anterior olfactory cortex, piriform cortex and taenia tecta. Whether olfactory region-projecting BF Npas1^+^ neurons also mediate arousal and motivational functions is an open question but conceivably, odors with strong survival/reproductive relevance could activate BF Npas1 neurons to enhance olfactory discrimination, arousal and appropriate motivational responses.

Previous chemogenetic studies had shown that activation of all GABAergic neurons in the BF (Anaclet et al., 2015) or in the VP subregion of BF (Li et al., 2021) strongly enhance wakefulness, whereas chemogenetic activation of vGlut2^+^ or cholinergic neurons at the same dose of CNO did not (Anaclet et al., 2015). However, these previous studies did not identify which of the many subtypes of BF GABAergic neurons (Yang et al., 2017) are responsible for this wake-promoting action. Although optogenetic excitation of the BF PV-expressing GABAergic subpopulation promotes arousals from sleep (Thankachan et al., 2019; McKenna et al., 2020) and cortical gamma band activity (Kim et al., 2015; Hwang et al., 2019; McNally et al., 2021), this effect was transient and only slightly increased total wakefulness, making it unlikely that activation of BF PV^+^ neurons is responsible for the strong wake-promoting effect of activating BF GABAergic neurons. Thus, our findings here represent a major advance in identifying a marker (Npas1) for strongly wake-promoting BF GABAergic neurons. Further analysis of our data revealed that the increase in wakefulness was due to an increase in long bouts of wakefulness and not due to more frequent awakenings. Concomitantly, the number of short NREM bouts was increased at the expense of long NREM bouts. Npas1^+^ neuron-mediated suppression of REM sleep was due to a reduced number of entries into REM. Control experiments in a separate group of mice confirmed that 0.3 mg/kg CNO administered without hM3d(Gq) transduction did not affect the amount of sleep or sleep architecture, consistent with other published findings that low doses of CNO do not have marked effects on sleep or sleep physiology (Anaclet et al., 2015; Traut et al., 2023). A previous study confirmed that CNO depolarizes and increases action potential discharge of BF Npas1^+^ neurons in the VP subregion transduced with hM3d(Gq)-mCherry (Morais-Silva et al., 2023), promoting activation of the immediate early gene product, cFOS. Here we confirmed that injections of the low dose of CNO (0.3 mg/kg) strongly increased the percentage of Npas1^+^ neurons expressing cFOS using IHC staining. Collectively these results strongly suggest that the behavioral results are due to activation of BF Npas1^+^ neurons. Furthermore, chemogenetic activation of Npas1+ neurons is associated with small but significant increases in the percentages of BF PV^+^ or ChAT^+^ neurons expressing c-Fos following CNO injections, likely via long-loop circuits and consistent with their wake-active profile (Xu et al., 2015).

Several potential targets of BF Npas1^+^ neurons may be involved in promotion of wakefulness. Optogenetic activation of VP GABAergic terminals in the VTA or lateral hypothalamus promotes wakefulness, whereas activation of terminals in the lateral habenula or mediodorsal thalamus does not (Li et al., 2021). Thus, the projections to the VTA and LH are strong candidate pathways to mediate the wake-promoting action. Additional projection sites which might mediate the wake-promoting actions of BF Npas1^+^ neurons are those to the supramammillary nucleus (Pedersen et al., 2017) and the shell of the nucleus accumbens (Luo et al., 2018). Overall, BF Npas1^+^ neurons appear to be part of subcortical circuits involved in motivationally driven wakefulness (Li et al., 2021). Consistent with that interpretation, chemogenetic activation of Npas1^+^ neurons increased EEG power in the high theta/alpha EEG band which has been linked to goal-directed behavior during wakefulness (Vassali & Franken, 2017; Fifel et al., 2022).

The functions of NREM sleep are linked to the specific cortical oscillations which define this state. Sleep spindles are linked to protection of sleep from the arousing effects of sensory stimuli and to sleep-dependent memory consolidation, especially when coupled to slow oscillations (Fernandez & Luthi, 2020). Slow oscillations in NREM sleep are linked to glymphatic clearance of toxic proteins, synaptic homeostasis and cellular metabolism. In our experiments, activation of Npas1^+^ neurons not only delayed the onset and amount of NREM sleep, but also reduced cortical slow wave and sigma band activity during NREM. Notably, the density, amplitude and duration of sleep spindles were significantly reduced, reminiscent of findings in various conditions such as schizophrenia, Alzheimer’s disease, and neurodevelopmental disorders (Fernandez & Luthi, 2020). Thus, the insomnia-like phenotype and disrupted NREM sleep oscillations produced by increased activity of BF Npas1^+^ neurons has face validity to human sleep and neuropsychiatric disorders.

Several studies have linked the Npas1 transcription factor and BF Npas1^+^ neurons to aspects of neuropsychiatric disorders and stress responsivity. Erbel-Sieler et al. (2004) found that Npas1 and Npas3 transcription factors are primarily expressed in inhibitory interneurons in cortical regions and, when knocked out, elicit behavioral and neurochemical abnormalities reminiscent of rodent models of neuropsychiatric disease such as schizophrenia and anxiety disorders. While abnormalities in cortical Npas1^+^ interneurons could mediate these behavioral abnormalities, a contribution of GPe or BF Npas1^+^ neurons cannot be ruled out. Michaelson et al., (2017) found that Npas1 is a master regulator of neuropsychiatric risk genes in hippocampal interneurons which are likely derived from the same ganglionic precursors which generate BF and GP Npas1^+^ GABAergic projection neurons. More recently, a role for VP Npas1^+^ neurons in stress responsivity has been demonstrated (Morais-Silva et al., 2023). Our study is the first to suggest a role for BF GABAergic neurons derived from the Npas1 lineage in sleep-wake control. Collectively, these findings suggest that overactivity of BF Npas1^+^ neurons could be important in stress-induced insomnia or sleep disruptions in neuropsychiatric disorders. Identification of this major population of wakefulness-promoting BF GABAergic neurons will allow new studies of their role in sleep disorders, dementia and other neuropsychiatric conditions affecting the BF. Furthermore, as far as we are aware, this is the first study to examine the sleep-wake function of a subgroup of BF neurons based on a developmental lineage marker, emphasizing the usefulness of developmental transcription factors in identifying novel subpopulations of sleep-wake control neurons (Liu et al., 2017; Kirjavainen et al., 2022).

## METHODS

### Animals

Npas1-cre-2A-tdTomato mice were provided by C. Savio Chan to establish a colony at VA Boston Healthcare System. The generation and validation of these mice, which are also available at Jackson Laboratory as strain 027718, has been previously described (Hernandez et al., 2015). Male hemizygous Npas1-cre-2A-tdTomato mice were crossed with female wild-type C57/BL6J mice (Jackson Laboratory strain 000664) mice to generate hemizygous mice for experiments. Genotype was verified by genotyping (Transnetyx, TN, USA). To identify BF vGlut1+ or vGlut2+ glutamatergic neurons, we crossed vGlut1-cre (Strain 023527; Jackson Laboratory) or vGlut2-cre mice (Congenic strain 028863; Jackson Laboratory) with a Cre-reporter strain expressing a red fluorescent marker (tdTomato; Strain 007905; Jackson Laboratory) to generate vGlut1-cre-tdTomato or vGlut2-cre-tdTomato mice respectively. The selective expression of Cre in vGlut1-cre mice has been validated previously (Harris et al., 2014). The selective expression of Cre-driven reporters in vGlut2 neurons when crossed with vGlut2-cre mice has also been validated in many brain areas (Krenzer et al., 2011; Vong et al., 2011), including BF (Anaclet et al., 2015; McKenna et al., 2021). To identify BF GABAergic neurons, we used heterozygous GAD67-GFP knock-in mice (Tamamaki et al., 2003; Brown et al., 2008; McKenna et al., 2013) from our in-house colony. Male, heterozygous GAD67-GFP mice on a Swiss-Webster background were crossed with female wild-type Swiss-Webster mice. To test if there was co-localization of NPAS1 in GABAergic neurons, we crossed Npas1-cre-2A-tdTomato animals with GAD67-GFP animals and performed immunohistochemical staining for NPAS1 in GAD67-GFP mice. Both male and female animals were used for anatomical and sleep-wake experiments. We did not observe any obvious sex differences. Thus, results from male and female animals were pooled.

Mice were housed under constant temperature and a 12:12 light:dark cycle (7AM:7PM), with food and water available *ad libitum*. All experiments conformed to U.S. Veterans Administration, Harvard University, and U.S. National Institutes of Health guidelines on the ethical use of animals. All measures were taken to minimize the number of animals used and their suffering and experiments were carried out in accordance with the National Institute of Health Guide for the Care and Use of Laboratory Animals (NIH Publications No. 80-23). Experimental procedures were approved by the Institutional Animal Care and Use Committee (IACUC) of the VA Boston Healthcare System.

### Target area within the BF

Our target area in the BF for neuronal characterization, anterograde tracing and chemogenetic experiments was the same as in our previous studies (McKenna et al., 2013, 2021; Yang et al., 2014). We focused on intermediate areas of BF (horizontal limb of the diagonal band (HDB), magnocellular preoptic nucleus (MCPO), substantia innominata (SI) and ventral pallidum (VP)), where previous studies in the rat (Rye et al., 1984; Gritti et al., 2003; Henny and Jones, 2008) and mouse (Anaclet et al., 2015; Do et al., 2016) found neurons projecting to the neocortex, approximately centered in BF at AP +0.14 mm; ML 1.6 mm; and DV −5.3mm. In contrast to our previous anatomical studies (McKenna et al., 2013, 2021) which were based on the Franklin and Paxinos (2008) brain atlas, we used semi-automated analyses based on the Allen Brain Atlas to facilitate throughput and comparison with other studies. This brain atlas combines the SI and VP subregions into one area and as such we use the term SI/VP to refer to this subregion of BF. As noted in the main text, there are also other subtle differences of the boundaries of BF subregions, when comparing the Allen Brain Atlas with that of Frankin and Paxinos (2008). Our analysis did not include the rostral aspect of BF (medial septum, vertical limb of the diagonal band (MS/DBV)), containing neurons largely projecting to the hippocampus, although Npas1^+^ neurons were also present in these areas. We also did not include the most caudal aspect of the cholinergic BF which extends into and intermingles with neurons in the GPe since the anatomy and physiology of Npas1^+^ neurons in the GPe have been extensively characterized in previous studies (Hernandez et al., 2015; Abecassis et al., 2020).

### General methods for anatomical studies

Mice were deeply anesthetized with sodium pentobarbital (50 mg/ml), exsanguinated with saline and perfused transcardially with a solution of 10% buffered formalin. Brains were post-fixed for 2 days in 10% formalin, and then transferred to a 30% sucrose solution for at least 24 hrs at 4°C. Tissue was cut at a thickness of 40 µm on a freezing microtome and collected into four wells of PBS. Following all immunohistochemistry procedures, tissue was washed and mounted onto gelatin-coated slides, dried, and coverslipped using Vectashield Vibrance antifade hard set mounting medium with DAPI, or without DAPI when blue secondary antibodies were used (H-1800 and H-1700; Vector Laboratories, Inc., Burlingame, CA).

### Immunohistochemistry methods

Prior to all immunohistochemical stains, coronal sections from one well were washed with PBS and placed in a blocking solution for one hour (0.5% TritonX-100 in 1X PBS + 3% normal donkey serum (NDS)). All primary antibody solutions were made in PBS with 0.3% Triton X-100 and 1% NDS, and secondary solutions in PBS and 1% NDS. The tissue was twice rinsed with phosphate buffered saline (5 mins/rinse) between primary and secondary antibody incubations. Details of the primary and secondary antibodies are provided in **Supplemental Tables 1 and 2**. Npas1-cre-2A-tdTomato mice express tdTomato constitutively but at low levels (Hernandez et al., 2015). Thus, anti-RFP staining was used to enhance the red signal. Rabbit primary anti-RFP (anti-DsRed Living Colors, 632496, Takara, Mountain View, CA, 1:1000) was followed with donkey anti-rabbit AF594 (715-586-152, Jackson, 1:100). To identify *neurons* expressing Npas1^+^ protein we used a previously described antibody (Hernandez et al., 2015) raised in guinea-pig (anti-NPAS1, 1:1000; validated in Npas1 knockout mouse tissue) followed by either donkey anti-guinea pig AF488 (706-545-148, Jackson, 1:500) or AF594 (706-585-148, Jackson, 1:500) for 2 hours at room temperature. To identify *cholinergic neurons* tissue was incubated in rabbit anti-choline acetyltransferase (ChAT, the synthetic enzyme for acetylcholine) for three days at 4°C (1:150; Cat#AB143, EMD Millipore, Billerica, MA) followed by incubation in a secondary antibody coupled to a green (donkey anti-rabbit AF488, Cat#A21206, Thermo Fisher Scientific, Cambridge, MA) fluorophore for 3 hrs at room temperature (1:200; RT). To identify *parvalbumin (PV)* neurons, tissue was incubated in sheep anti-PV overnight at 4°C (1:150; Cat#AF5058, RnD Systems, Minneapolis, Mn) followed by a secondary antibody (1:200) coupled to a green (donkey anti-sheep AF488 Cat#A11015, Thermo Fisher Scientific) fluorophore for 3 hrs at RT. To *amplify EYFP* signal for anterograde tracing using AAV5-DIO-ChR2-EYFP, tissue was incubated in mouse polyclonal anti-GFP primary antibody (1:300; Cat#MAB3580, EMD Millipore) for 3 days at 4**°**C, and secondary antibody (1:500; donkey anti-mouse AF488, Cat#A21202, Thermo Fisher Scientific) overnight (16 hrs) at 4°C. To identify neurons which were activated during chemogenetic experiments where Npas1^+^ neurons were identified by the presence of mCherry (red) we performed immunohistochemical staining for the immediate early gene cFOS, using a primary antibody raised in rabbit (1:200 overnight at 4°C; EMD Millipore Cat#ABE457) followed by donkey anti-rabbit secondary antibody (1:200 for 2hrs at RT; Thermofisher Cat#A-21206;) conjugated to a green fluorophore (AF488). To identify ChAT^+^ or PV^+^ neurons in tissue with mCherry (red) and cFos stained-AF488 (green) neurons/nuclei, we incubated the tissue with the primary antibodies against ChAT and PV described above with a blue secondary antibody (donkey anti-rabbit AF350, Cat#A10039, Thermo Fisher Scientific; 1:200) or (donkey anti-sheep AF350, 1:200, Cat#A21097, Thermo Fisher Scientific) for 3 hrs at room temperature. All primary (**Supplemental Table 1**) and secondary (**Supplemental Table 2**) antibodies used here have been previously validated and used in peer reviewed publications. Secondary antibody-only controls were also performed, omitting the above-listed primary antibodies.

### AAV5-DIO-ChR2-EYFP injection for anterograde tracing of the projections of BF Npas1^+^ neurons

#### Viral vectors

Viral vectors encoding Channelrhodopsin-2 (ChR2) and enhanced yellow fluorescent fusion proteins (AAV5-DIO-ChR2-EYFP) originally generated by the laboratory of Dr. Karl Deisseroth (Stanford University, CA, USA) were purchased from the University of North Carolina Vector Core facility.

#### Stereotaxic surgery and viral injection

Mice were deeply anesthetized with isoflurane (1-3%) and viral injections were performed using a 1 µl Hamilton syringe (Cat#87900, Point Style 2, Hamilton Company, Reno, NV, USA), targeting BF (AP +0.14 mm; ML 1.6 mm; DV −5.3mm). 300 nl of viral vector was injected unilaterally at a flow rate of 50 nl/min. The needle was left in place for an additional 10 mins to allow virus diffusion in the brain and to avoid backflow along the needle tract. Once viral injections were complete, the needle was slowly removed. Animals were sacrificed following one month of survival post-injection.

### Chromophore co-localization quantification/photographic depiction

For each fluorophore, representative BF sections (three 40 µm coronal sections per animal, spanning +0.38 mm (rostral), +.014 mm (medial), and −0.10 mm (caudal) from bregma)) were imaged using StereoInvestigator and quantified using NeuroInfo software packages (Version 2022; Microbrightfield, Williston, VT) and a Zeiss Imager.M2 fluorescent microscope. Neurons labeled with tdTomato or RFP were identified by the presence of red (excitation:emission at 590:617 nm) fluorescence in the cytoplasm and nucleus. Neurons labeled with GFP/green secondary antibody or a blue secondary antibody were identified by were identified by the presence of green or blue (excitation:emission 488:509 nm and 343:441 nm, respectively) fluorescence in the cytoplasm (PV or ChAT stains) or nucleus (cFOS or NPAS1 IHC stains). Green staining was observed in the cytoplasm and nucleus for GAD67-GFP mice, as in our previous studies (McKenna et al., 2013; 2021; Yang et al., 2013, 2014). There was a high background level of GFP in the neuropil due to the presence of GFP in GABAergic fibers, particularly in the dorsal BF. In our previous studies using these mice (McKenna et al., 2013; 2021), non-GABAergic neurons, i.e. ChAT^+^ or vGlut2^+^ neurons, demonstrated GFP fluorescence which was below background levels, and we used the same criterion to identify GFP-negative nuclei here.

The BF region of each section was roughly traced under low (5X) magnification. The 3D Slide Scanning Workflow feature of StereoInvestigator was then used to capture and stitch together individual Z-stacks taken within the outlined BF region at a higher (10X) magnification. Each stack was comprised of 8 focal planes between the top and bottom of the tissue slice, manually determined at upper and lower stage height boundaries where clear focus is lost. Once a composite Z-stack of the entire BF was taken, the image was then collapsed to a single plane using the Deep Focus feature in NeuroInfo to visualize cells and nuclei throughout the depth of the section. An initial pass at quantification was automatically performed using the Cell Detection function of NeuroInfo, which identifies fluorescent puncta from size and intensity. To reduce false positives, diameters were conservatively set between 5-7 µm for nuclear stains and 10-15 uM for cytoplasmic stains. Automatically detected cells were manually confirmed and additional quantification within the BF was manually performed as needed for additional undetected neurons. Colocalization was first assessed automatically by distance (≤7 µm between labeled puncta) and confirmed by manual inspection.

After cells in the BF were labeled, the Section Registration function of NeuroInfo was used to align the captured image to its corresponding plane in the 3D Allen Mouse Brain Atlas, correcting for the sectioning angle and any skew in X and Y dimensions resulting from mounting or post-fixation distortion. Regions of interest (SI/VP, HDB, MCPO, and surrounding areas) were plotted onto the section image, and the total number of marked cells in each region was then exported. Cell density was calculated by dividing the total number of cells in each subregion by its area in µm^2^.

#### Long-axis measurement

The long-axis diameter of Npas1-tdTomato labeled neurons was measured for each of the 3 BF subnuclei using NeuroInfo software and images collected as described above. 10 neurons per BF subnuclei (HDB, MCPO, SI/VP, 5 neurons/hemisphere) were randomly selected and measured for the 3 representative sections (rostral, intermediate, and caudal) in each animal.

#### Anterograde tracing

EYFP fluorescence in mice injected with AAV5-DIO-ChR2-EYFP was first amplified with anti-GFP antibody and secondary antibody conjugated to a green fluorophore. 40 µm coronal sections were mounted, incorporating tissue from the most rostral to most caudal sections and spaced 80 µm apart, anatomically. Whole-slide scans were performed at 20x magnification utilizing the PhenoImager HT automated multispectral imaging system (Akoya Bioscience, Marlborough, MA). BF Npas1^+^ efferent regions (AAV-EYFP anterograde projections) were identified as regions where varicosities of AAV-EYFP-labeled axons were located. The relative abundance of individual projections was agreed upon by 2 or more experimenters.

### Chemogenetics and stereotaxic surgery

To test the effect of activation of BF Npas1^+^ neurons on sleep-wake behavior, adeno-associated viral vectors expressing excitatory receptors and a fluorescent marker of expression (pAAV-hSyn-DIO-hM3D(Gq)-mCherry, plasmid #44361, Addgene, Watertown, MA) were injected bilaterally into BF (AP: + 0.4, ML: ± 1.6, DV: −5.3) at 0.5, 0.7 or 1 µl/side of adult (3-6 month old) mice. The number of neurons transduced and sleep-wake effects were comparable among the different doses and have been pooled. To record cortical electrical activity, bilateral frontal neocortical EEG screw electrodes (Pinnacle Technology Inc.; Kansas, United States; Part # 8403) were placed at AP: +1.5 mm, ML: ± 0.5 with a ground electrode at AP: −3 mm, ML: +2.7 and a reference electrode at AP: −6 mm ML: 0, and soldered to a headmount (Pinnacle Technology Inc.; Part # 8201-SS). EMG electrodes were placed in the nuchal muscle. All chronically implanted components were secured with dental cement (Keystone industries, Bosworth Fastray; Part # 0921378). Four or more weeks later (to allow for viral transduction), sleep-wake recordings began at 7AM (ZT0). Intraperitoneal injections of saline (vehicle control) or 0.3 or 1 mg/kg CNO were given early in the light cycle at 9AM (ZT2) when mice normally sleep, as we predicted an increase in wakefulness based on previous results investigating all BF GABAergic neurons (Anaclet et al., 2015). Control CNO injection experiments were conducted in a different strain of transgenic mice (Lhx6-cre) on the same C57/BL6J background strain of mice using a control virus expressing a fluorescent marker without hM3d(Gq) (AAV-DIO-EYFP). Following completion of sleep-wake recordings and 120 min prior to sacrifice, mice received an additional injection of saline or CNO (0.3 or 1 mg/kg) at ZT2 verify activation of Npas1^+^ neurons using immunohistochemical staining for the immediate early gene cFOS.

### Sleep-wake recordings and analysis

Mice were tethered and acclimatized to the recording chamber for at least 3 days. After acclimatization, continuous EEG/EMG (8200-K1-SL amplifier; Pinnacle Technology Inc., Lawrence, KS, USA) recordings were performed before, during, and after i.p. injections of CNO or saline (vehicle control) for 24 hrs from ZT0-24. EEG/EMG were recorded using Sirenia Sleep Pro (Pinnacle Technology). EEG signals were sampled at 1 KHz, amplified 100x and bandpass filtered at 0–500 Hz. EMG signals were filtered at 0-100 Hz. As described previously (McKenna et al., 2020; Uygun et al., 2022) sleep-wake states were manually scored in 4 s epochs as follows: Wake was scored when EEG showed a desynchronized low amplitude signal with muscle tone evident by a large EMG signal; the large EMG signal did not need to be phasic in appearance to be characterized as wake. NREM sleep was scored when the EEG signal showed large amplitude, slow synchronized waves, and a low EMG signal, except for very brief bursts which were considered twitching. REM sleep was scored when the EEG signal presented a repetitive stereotyped ‘sawtooth’ signal in the theta range (4-9 Hz), with a nearly flat EMG signal. Artifacts were dealt with as follows: periods that appeared to be wakefulness, but the EEG was contaminated by crosstalk with EMG signals, were scored as ‘wake-exclude’. Periods of NREM or REM with large amplitude DC shifts were extremely rare and labelled as ‘NREM-exclude’ or ‘REM-exclude.’ All scored epochs were used in behavioral analyses such as time in state or bout analysis, but epochs with artifacts were excluded for analyses of EEG signals, such as power spectral density or time-frequency analyses. Scoring was performed in Sirenia Sleep, and EEG signals and scored epochs were exported for further analysis in MATLAB.

Power spectral density of wake, NREM and REM was computed using the MATLAB pwelch function using a 4-second Hanning window with 50% overlap. Frequency bands were defined as follows: slow wave: 0-1.5 Hz; delta: 1.5-4 Hz; theta: 4-9 Hz; alpha/sigma: 9-15 Hz; beta: 15-30 Hz; gamma: 30-80 Hz. To normalize EEG power, power in each of these frequency bands per state was divided by the total power across all frequencies (0-500Hz) from the entire 24 hr record. Power spectra were binned in 3-hour intervals, except for the NREM delta power timecourse throughout the light cycle (**Supplemental Figure 5B**), which used 1-hour bins. Normalized power per state per band was averaged within treatment groups. We analyzed sleep spindles (≥0.5 s in length) using our validated automated spindle detection algorithm (Uygun et al., 2019), where normalized amplitude is calculated by dividing against a threshold value based on putative spindle peaks in each EEG record. Representative time-frequency spectrograms were created using the mtspectrogramc function (Chronux Toolbox; Chronux.org).

Bout durations were binned into discrete intervals, summed, and presented as a percentage of total time in the state. Latency to NREM sleep was defined as the time post-injection until the start of 10 continuous NREM epochs (40 sec). Latency to REM sleep was defined as the time post-injection until the start of 5 continuous REM epochs (20 sec). Brief awakenings were defined as a ≤ 4 epoch (≤ 16 sec) long period of wake interrupting 2 NREM epochs.

### Statistical Analysis

Data are presented as mean ± SEM. All neuroanatomical density, long-axis diameter, co-localization measures, and comparisons of group averages (i.e. average sleep latencies or spindle densities) were analyzed using one-way analysis of variance (ANOVA), to compare differences between BF subnuclei (HDB, MCPO, SI/VP), coronal BF section representations (rostral, medial, and caudal), or drug treatments. Two-way ANOVAs were performed to analyze treatment effects over time. When appropriate in an individual ANOVA analysis, repeated-measures (RM) factor-matching was performed. If a main effect of the ANOVA was significant, pair-wise comparisons between groups were then made, using Holm-Šídák multiple comparison corrections. NREM spindle analyses over time were performed using two-way mixed-effect models to facilitate RM two-factor analyses when individual datapoints within subjects were excluded (i.e. a mouse exhibited no NREM sleep during a particular bin). Comparisons between 2 group means were performed as two-tailed t-tests and paired when appropriate. Cumulative histograms are compared as two-sample Kolmogorov-Smirnov (K-S) tests. Statistics were performed in GraphPad Prism (Version 9.X and 10.X, GraphPad Software, Boston, MA), MATLAB (Version 2023a, The MathWorks Inc, Natick, MA) or Microsoft Excel (Version 2016, Microsoft, Redmond, WA).

## DECLARATIONS

### Availability of data

Portions of these results have been previously reported in abstract form (Yang et al., 2019; Brown et al., 2021; Troppoli et al., 2023). Data will be uploaded to a public repository upon acceptance.

### Code availability

The MATLAB script for spindle analysis was described in Uygun et al., 2019 and is available upon request.

### Authors’ contributions (for discussion, edit as needed)

T.A.T. performed immunohistochemical and chemogenetic experiments in Npas1-cre-2A-tdTomato mice, conducted anatomical mapping of projections, analyzed the data and edited the manuscript. C.Y. performed a subset of the chemogenetic experiments. F.K. performed a subset of the surgeries and perfusions for the anatomical tracing experiments. D.S.U. provided guidance on analysis of sleep-wake data. I.L., D.A., and T.S. performed immunohistochemical staining, transcardial perfusions and/or tissue mounting. J.M.M. provided expert technical guidance and comments on the manuscript. R.B. provided funds and comments on the manuscript. C.S.C. generated the Npas1-cre-2A-tdTomato mice and NPAS1 antibodies and edited the manuscript. J.T.M. performed, oversaw, designed and analyzed anatomical experiments to characterize Npas1^+^ neurons in the basal forebrain of Npas1-cre-2A-tdTomato mice and edited the manuscript. R.E.B. obtained funding, conceived the study, designed experiments, interpreted data and wrote and edited the manuscript.

### Ethics approval

All experiments were reviewed and approved by the institutional animal care and use, safety, biosafety and research and development committees of VA Boston Healthcare System.

### Consent to participate

N/A.

### Consent for publication

All authors approved this submission.

## Table of abbreviations

3V: Third ventricle
AAV: Adeno-associated virus
Aco: Anterior commissure, olfactory limb
Act: Anterior commissure, temporal limb
ADP: Anterodorsal preoptic nucleus
AI: Agranular insular cortex AMY Amygdala
BF: Basal forebrain
BNST: Bed nucleus of stria terminalis
CA1: Hippocampal field CA1
CPu: Caudate Putamen
cc: Corpus callosum
Cg: Cingulate cortex
Cg1: Cingulate cortex, area 1
ChAT: Choline acetyl transferase
ChR2: Channelrhodopsin 2
Cl: Claustrum
cp: Cerebral peduncle
CTX: Cortex
DBv: Vertical limb of the diagonal band
DEn: Endopiriform nucleus, dorsal
DG: Dentate gyrus
DM: Dorsomedial hypothalamic nucleus
DP: Dorsal peduncular cortex
DR: Dorsal raphe nucleus
EYFP: Enhanced yellow fluorescent protein
fx: Fornix
fmi: Forceps minor of the corpus callosum
fr: Fasciculus retroflexus
GAD67: Glutamic acid decarboxylase-67 kDa isoform
GFP: Green fluorescent protein
GPe: Globus pallidus, pars externa
HDB: horizontal limb of the diagonal band
ic: Internal capsule
EP: Endopiriform cortex, intermediate
IF: Interfascicular nucleus
IL: Infralimbic cortex
IMD: Intermediodorsal thalamic nucleus
LH: Lateral hypothalamic area
IPR: Interpeduncular nucleus, rostral
LDT: Laterodorsal tegmental nucleus
LHA: Lateral hypothalamic area
LHb: Lateral habenula
LM: Lateral mammillary nucleus
LO: Lateral orbital cortex
LP: Lateral posterior thalamic nucleus
LPO: Lateral preoptic area
LS: Lateral septum
LSUM: Supramammillary nucleus, lateral
M1: Primary motor cortex
M2: Secondary motor cortex
MCPO: Magnocellular preoptic nucleus
MD: Mediodorsal thalamic nucleus
Me: Medial amygdaloid nucleus
MHb: Medial habenula
ml: medial lemniscus
ML: Medial mammillary nucleus, lateral
MM: Medial mammillary nucleus, medial
MnPO: Median preoptic nucleus
MO: Medial orbital cortex
mp: Mammillary peduncle
MPA: Medial preoptic area
mPFC: Medial prefrontal cortex
MR: Medial raphe nucleus
MS: Medial septum
mt: Mammillothalamic tract
mtg: Mammillotegmental tract
NAc: Nucleus accumbens
ns: Nigrostriatal bundle
opt: Optic tract
PAG: Periaqueductal gray
PB: Parabrachial nucleus
PFC: Prefrontal cortex
Pir: Piriform cortex
pm: Principal mammillary tract
PN: Paranigral nucleus
Po: Posterior thalamic nuclear group
PrL: Prelimbic cortex
PV: Parvalbumin
PVT: Paraventricular thalamic nucleus
RE: Nucleus reuniens
RLi: Rostral linear raphe nucleus
RPC: Red nucleus, parvicellular
SEM: Standard error of the mean
SH: Septohippocampal nucleus
SI: Substantia innominata
SNC: Substantia nigra, compacta
SNR: Substantia nigra, reticulata
SUM: Supramammillary nucleus
TT: Tenia tecta
Tu: Olfactory tubercle
vGlut2: Vesicular glutamate transporter 2
VLPO: Ventrolateral preoptic nucleus
VM: Ventromedial thalamic nucleus
VMH: Ventromedial hypothalamic nucleus
VO: Ventral orbital cortex
VP: Ventral pallidum
VPT: Ventral posterior thalamic nuclei
VTA: Ventral tegmental area of Tsai
VTM: Ventral tuberomammillary nucleus
ZI: Zona incerta

