## Supplemental Figure 1 for "Neuronal PAS domain 1 identifies a major subpopulation of wakefulness-promoting GABAergic neurons in basal forebrain"

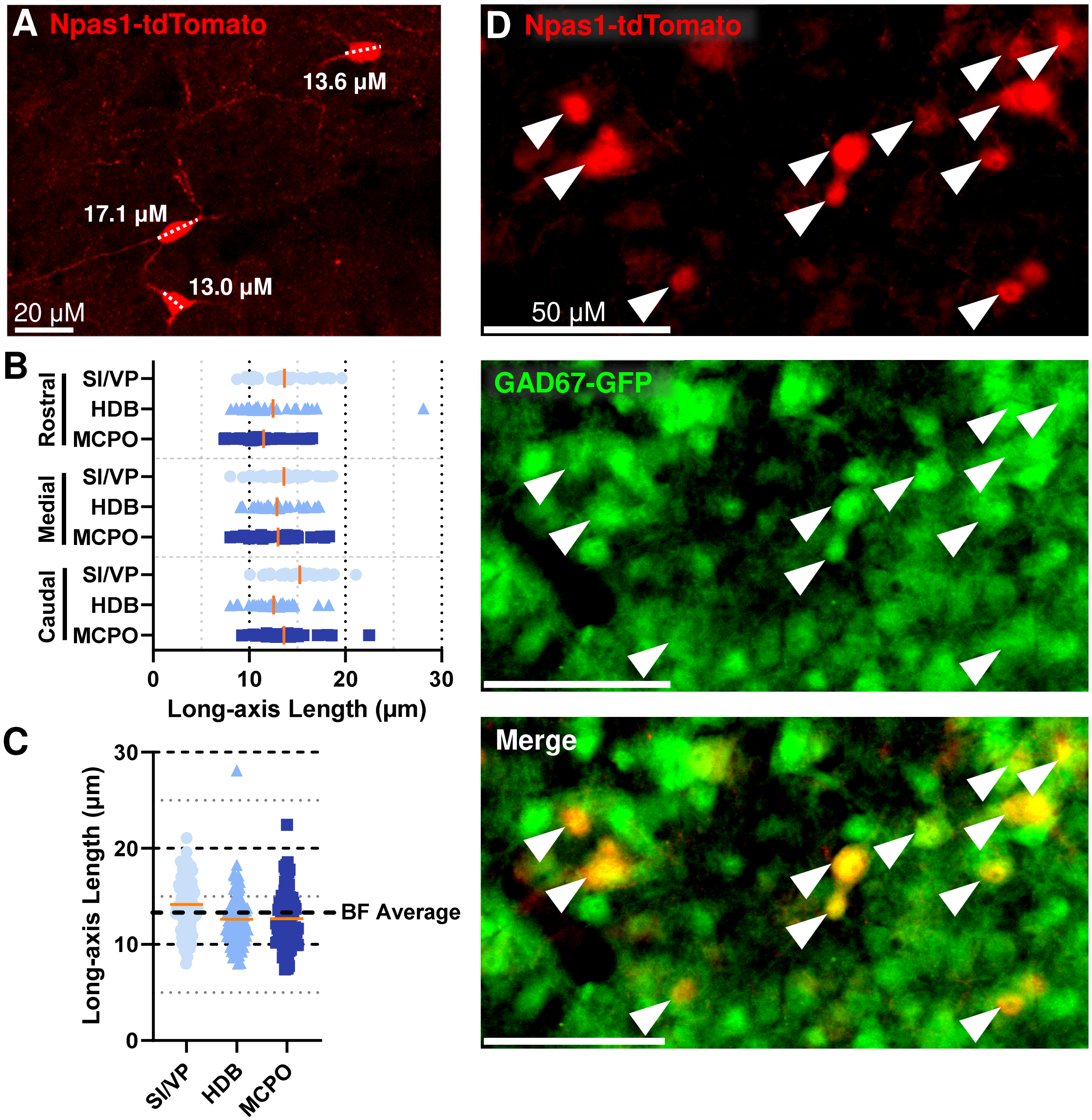

A Npas1-tdTomato
13.6 µM
17.1 µM
13.0 µM
20 µM
B
Rostral
Medial
Caudal
SI/VP
HDB
MCPO
0
10
20
30
Long-axis Length (µm)
C
Long-axis Length (µm)
BF Average
SI/VP
HDB
MCPO
D Npas1-tdTomato
50 µM
GAD67-GFP
Merge

**Supplemental Figure 1: Npas1$^+$ cells are small or medium sized neurons and colocalize with GAD67-expressing cells in a GAD67-GFP/Npas1-cre-2A-tdTomato mouse genetic cross. (A)** Example subset of BF Npas1-tdTomato neuron long-axis measurements. **(B)** Distribution of Npas1-tdTomato long-axis measurements from a random selection of 10 cells per BF subregion across rostral, medial, and caudal BF sections collected from each of 3 animals (270 cells total). Mean long-axis measurement per subregion is marked with an orange line. **(C)** Npas1-tdTomato cells were significantly longer in rostral vs. ventral BF subregions (SI/VP: 14.17 ± 0.35 µm, vs. HDB: 12.63 ± 0.55 µm, p=0.0010; MCPO: 12.70 ± 0.39 µm, p=0.0012, mean marked by an orange line). The overall length of Npas1-tdTomato cells in the BF was 13.2 ± 0.4 µm, indicated by a dashed black line. Comparisons made via one-way ANOVA ($F_{2,267}$ = 8.48, p<0.0001). **(D)** Example image within the BF showing colocalization of Npas1-tdTomato cells (red) with GAD67-GFP cells (green) following a GAD67-GFP/Npas1-cre-2A-tdTomato mouse cross. Colocalization is indicated with an arrowhead.
