## Supplemental Figure 2 for "Neuronal PAS domain 1 identifies a major subpopulation of wakefulness-promoting GABAergic neurons in basal forebrain"

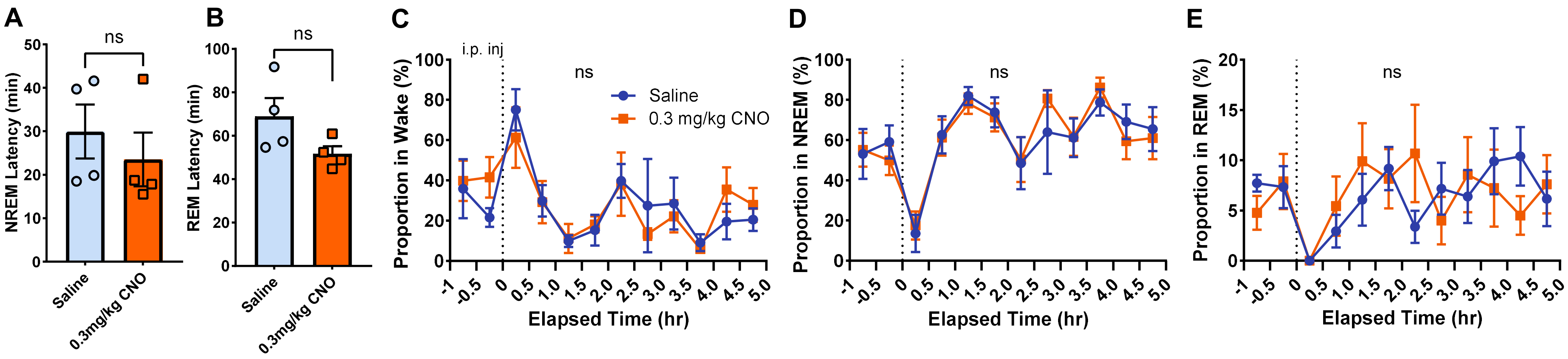

A
ns
NREM Latency (min)
Saline
0.3mg/kg CNO
B
ns
REM Latency (min)
Saline
0.3mg/kg CNO
C
i.p. inj
ns
Saline
0.3 mg/kg CNO
Proportion in Wake (%)
Elapsed Time (hr)
D
ns
Proportion in NREM (%)
Elapsed Time (hr)
E
ns
Proportion in REM (%)
Elapsed Time (hr)

**Supplemental Figure 2: CNO alone has no impact on sleep behavior in mice lacking pAAV-hSyn-DIO-hM3D(Gq). (A)** Latency to NREM sleep is unchanged between saline and 0.3 mg/kg CNO administration (n=4, p=0.28, two-tailed paired t-test). **(B)** Latency to REM sleep is also not significantly different between saline and 0.3 mg/kg CNO administration (n=4, p=0.17, two-tailed paired t-test). **(C)** The proportion of time spent awake was not significantly different between saline and 0.3 mg/kg CNO treated animals at any time point. Two-way ANOVA identifies no interaction between time x treatment ($F_{11,72}$ = 0.507 p=0.89). **(D)** Time spent in NREM sleep did not significantly differ between saline and 0.3 mg/kg CNO treated animals at any time point. Two-way ANOVA identifies no interaction between time x treatment ($F_{11,72}$ = 0.291 p=0.99). **(E)** The proportion of mice time spent in REM sleep was not significantly different between saline and 0.3 mg/kg CNO groups at any time point. Two-way ANOVA identifies no interaction between time x treatment ($F_{11,72}$ = 0.892 p=0.55). ns: not significant.
