## Supplemental Figure 3 for "Neuronal PAS domain 1 identifies a major subpopulation of wakefulness-promoting GABAergic neurons in basal forebrain"

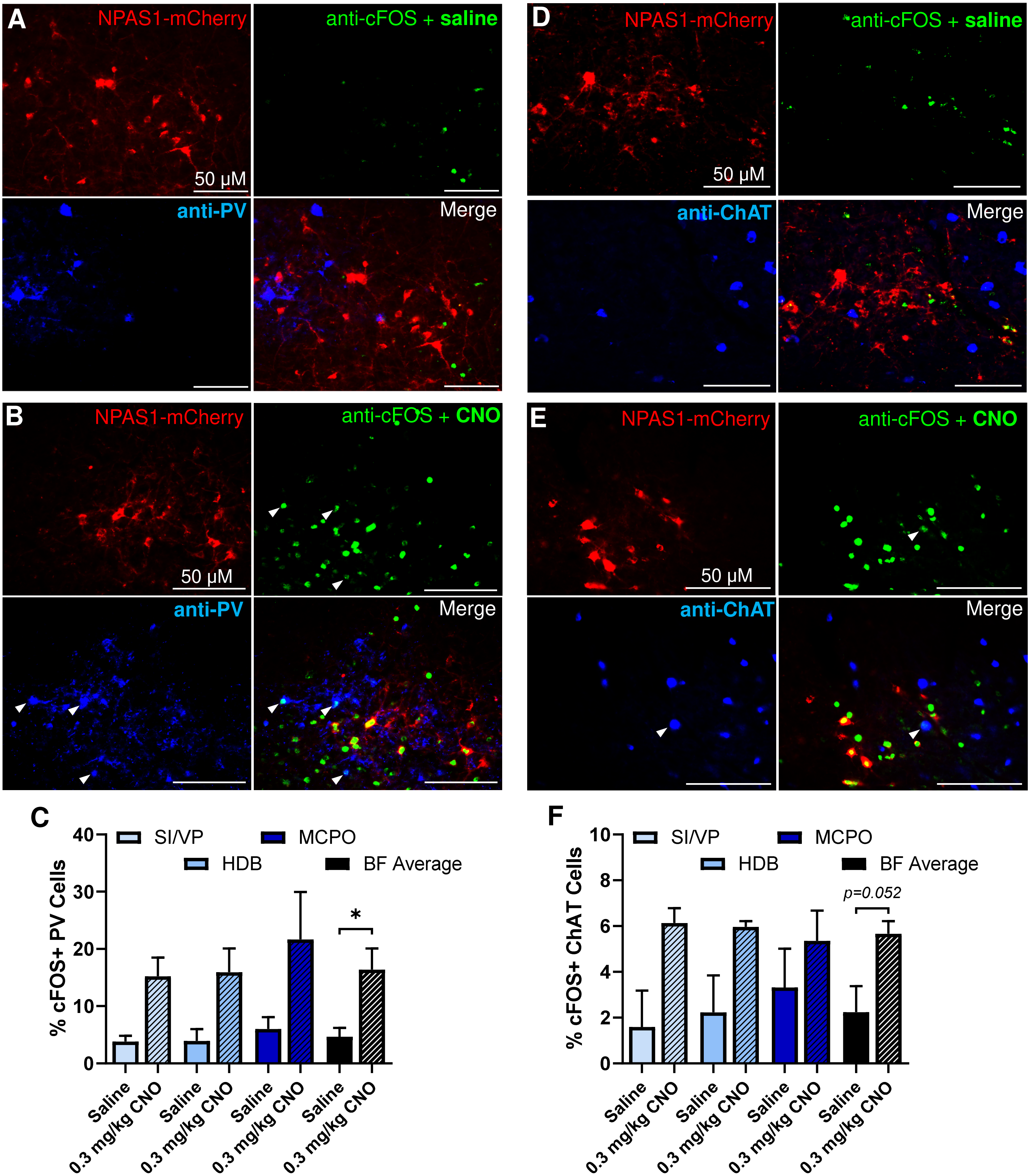
A
NPAS1-mCherry
anti-cFOS + saline
50 μM
anti-PV
Merge
B
NPAS1-mCherry
anti-cFOS + CNO
50 μM
anti-PV
Merge
C
SI/VP
MCPO
HDB
BF Average
% cFOS+ PV Cells
0
10
20
30
40
*
Saline
0.3 mg/kg CNO
D
NPAS1-mCherry
anti-cFOS + saline
50 μM
anti-ChAT
Merge
E
NPAS1-mCherry
anti-cFOS + CNO
50 μM
anti-ChAT
Merge
F
SI/VP
MCPO
HDB
BF Average
% cFOS+ ChAT Cells
0
2
4
6
8
10
p=0.052
Saline
0.3 mg/kg CNO

**Supplemental Figure 3: Chemogenetic activation of BF Npas1$^+$ cells activates PV and ChAT cells consistent with a wake-active profile. (A, B)** Representative immunohistochemically stained images depicting transduced Npas1-mCherry cells (red), an anti-cFOS nuclear stain (green), and a stain against PV cells (blue) collected from animals administered saline **(A)** or 0.3 mg/kg CNO **(B)** 2h prior to sacrifice. Colocalized cFOS and PV puncta are labeled with an arrowhead. **(D, E)** Representative images depicting transfected NPAS1-mCherry neurons (red), an anti-cFOS nuclear stain (green), and a stain against ChAT cells (blue) in BF tissue from animals given saline **(D)** or 0.3 mg/kg CNO **(E)** prior to sacrifice. C-FOS and ChAT colocalization is marked with an arrowhead. **(C)** Chemogenetic activation of BF Npas1$^+$ neurons with 0.3 mg/kg CNO (n=3 mice) increases wakefulness and significantly increased double-labeling of all BF PV cells with an anti-cFOS nuclear stain compared to saline administration (n=3; 4.65% ± 1.53% vs. 16.4% ± 3.72%, p=0.044; unpaired t-test). **(F)** The activation of BF Npas1$^+$ neurons and increased wakefulness following administration of 0.3 mg/kg CNO increases anti-cFOS and ChAT double-labeling (n=3) following saline administration (n=3; 2.24% ± 1.13% vs. 5.66% ± 0.55%, p=0.052; unpaired t-test). $p<0.05$ *.
