## Supplemental Figure 4 for "Neuronal PAS domain 1 identifies a major subpopulation of wakefulness-promoting GABAergic neurons in basal forebrain"

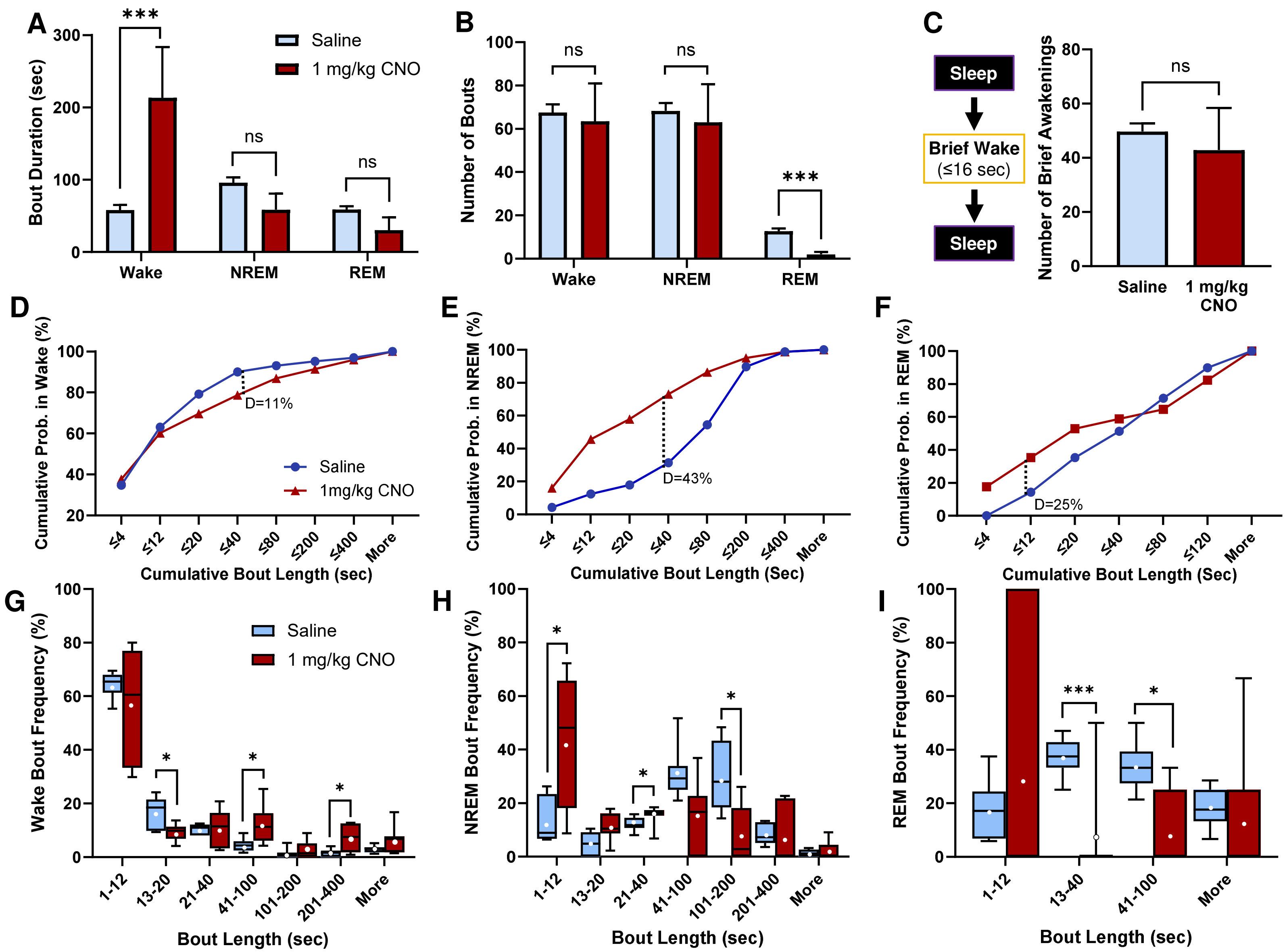
A
B
C
D
E
F
G
H
I
Bout Duration (sec)
Saline
1 mg/kg CNO
Wake
NREM
REM
ns
***
Number of Bouts
Sleep
Brief Wake
(≤16 sec)
Number of Brief Awakenings
Cumulative Prob. in Wake (%)
Cumulative Prob. in NREM (%)
Cumulative Prob. in REM (%)
1mg/kg CNO
D=11%
D=43%
D=25%
Cumulative Bout Length (sec)
Cumulative Bout Length (Sec)
Wake Bout Frequency (%)
NREM Bout Frequency (%)
REM Bout Frequency (%)
Bout Length (sec)

**Supplemental Figure 4: Chemogenetic activation of BF Npas1$^{+}$ neurons with 1 mg/kg CNO significantly lengthens wake bouts and suppresses the frequency of REM bouts. (A)** Compared to saline (n=7 mice), 1 mg/kg CNO (n=7) administration promotes significantly longer wake bouts over the 3-hour post-injection period (58.0 ± 7.1 sec vs 213.2 ± 70.52 sec respectively, p=0.0039; two-tailed paired t-test). Average NREM bout length was not significantly different following administration of saline compared to 1 mg/kg CNO (95.9 ± 7.6 sec vs 58.5 ± 22.2 sec respectively, p=0.14), nor was REM bout length (58.9 ± 4.3 sec vs. 30.0 ± 18.2, p=0.15). **(B)** The number of wake bouts was not significantly different between saline and 1 mg/kg CNO administration during the 3-hour post-injection period (67.6 ± 3.7 vs. 63.4 ± 17.6 respectively, p=0.85), nor were the number of NREM bouts (68.3 ± 3.7 vs. 63.0 ± 17.6, p=0.80). However, compared to saline, 1 mg/kg CNO administration produced significantly fewer REM bouts (12.7 ± 1.3 vs. 2.0 ± 1.1, p=0.00056. **(C)** The number of brief awakenings (≤16 sec of wake interspersing 2 sleep bouts) was not significantly different between saline and 1 mg/kg CNO treatments (49.7 ± 3.0 vs. 42.9 ± 15.5, p=0.70). **(D)** A cumulative probability plot of wake bout lengths during the 3-hour post-injection period indicates 1 mg/kg CNO produces significantly less frequent short wake bouts (p=0.0031, K-S D=0.11). **(G)** Bout analysis finds significantly fewer short wake bouts between 13-20 sec (p=0.045), and significantly more between 41-100 sec (p=0.039), and 201-400 sec (p=0.019) following 1 mg/kg CNO administration vs. saline. **(E)** NREM bout length cumulative distributions are significantly different between saline and 1 mg/kg CNO administrations, favoring shorter NREM bouts (p<0.0001, D=0.43). **(H)** 1 mg/kg CNO produces significantly more short-duration NREM bouts between 1-12 sec (p=0.025) and 21-40 sec (p=0.024), and significantly fewer long bouts between 101-200 sec (p=0.019). **(F)** Cumulative distributions of REM sleep are not significantly different between treatments (p=0.30, K-S D=0.25). **(I)** 1 mg/kg CNO administration produces significantly fewer REM bouts between 13-40 sec (p=0.0041) and 41-100 sec (p=0.039). p<0.05 *, p<0.005 ***. ns: not significant. Cumulative histograms are compared as two-sample nonparametric Kolmogorov-Smirnov (K-S) tests, and individual binned bout comparisons were made via two-tailed paired t-tests. Whiskers on box and whisker plots range from min to max, the midline represents the median, and the mean is indicated with a white circle. Up to 1 animal per group bin may be excluded if it is alone in spending 0% or 100% of its time in that bin. p<0.05 *, p<0.01 **, p<0.005 ***, ns: not significant.
