## Supplemental Figure 5 for "Neuronal PAS domain 1 identifies a major subpopulation of wakefulness-promoting GABAergic neurons in basal forebrain"

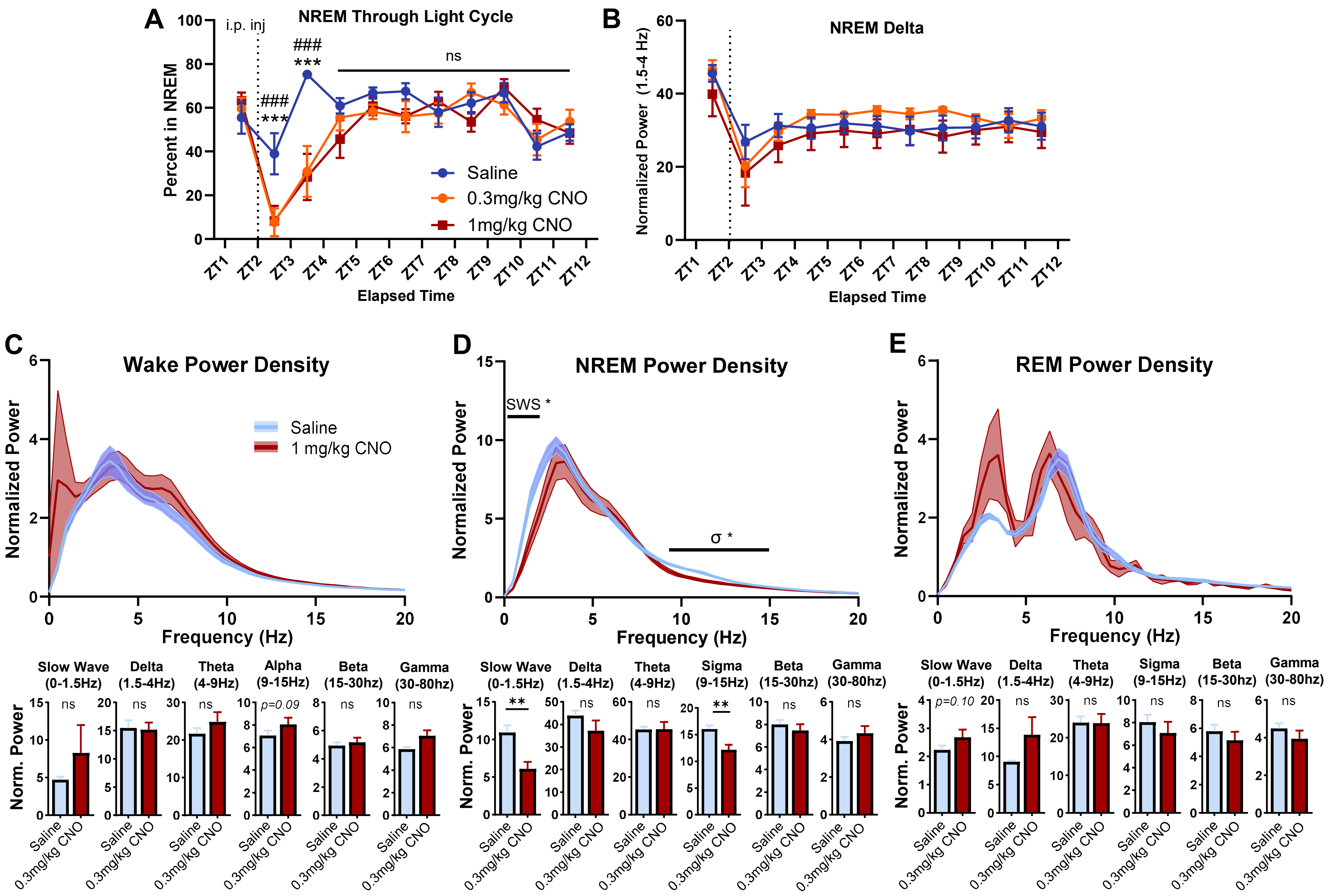

A
NREM Through Light Cycle
i.p. inj
Percent in NREM
Elapsed Time
Saline
0.3mg/kg CNO
1mg/kg CNO
###
***
ns
B
NREM Delta
Normalized Power (1.5-4 Hz)
Elapsed Time
C
Wake Power Density
Normalized Power
Frequency (Hz)
Saline
1 mg/kg CNO
D
NREM Power Density
SWS *
σ *
E
REM Power Density
Slow Wave (0-1.5Hz)
Delta (1.5-4Hz)
Theta (4-9Hz)
Alpha (9-15Hz)
Sigma (9-15Hz)
Beta (15-30hz)
Gamma (30-80hz)
Norm. Power
p=0.09
p=0.10
**
Saline
0.3mg/kg CNO

**Supplemental Figure 5: CNO administration and Npas1$^{+}$ activation does not result in either NREM or delta power rebound during the light period, but 1 mg/kg CNO does significantly alter NREM spectral EEG activity. (A)** Administration of 0.3 mg/kg (n=9) and 1 mg/kg CNO (n=7) suppresses NREM sleep compared to saline (n=9) for 2 hours post-injection. Afterwards, mice spend similar proportions of time in NREM throughout the duration of the light cycle. Two-way ANVOA identifies significant interaction of time x treatment ($F_{20,198} = 3.0$, $p<0.0001$). **(B)** There is no significant difference in NREM delta power (1.5-4Hz) across treatments at any timepoint. Two-way ANVOA identifies no interaction of time x treatment ($F_{20,171} = 0.91$, $p=0.58$). **(C)** A normalized power density plot and power spectra analyses identifies a trend for higher wake alpha power (9-15Hz, maximally between 11.7-12.2Hz) following 1 mg/kg CNO (n=7) vs saline (n=7) administrations (p=0.090, comparisons made via two-tailed paired t-tests). No significant changes were observed in slow wave (p=0.36), delta (p=0.79), theta (p=0.17), beta (p=0.44) or gamma power bands (p=0.35). **(D)** Power density spectra and normalized power band analysis indicates that 1 mg/kg CNO administration produces significant less power in the slow wave (0-1.5Hz, maximally at 0.49-0.98Hz, p=0.0086) and sigma power bands (9-15Hz, maximally at 10.7-11.2Hz, p=0.0094). No significant differences were seen in delta (p=0.19), theta (p=0.99), beta (p=0.44) or gamma bands (p=0.31). **(E)** Frequency spectra were not significantly different between treatments during REM sleep, though there was a trend toward increased slow-wave activity (p=0.099). Other bands showed no change, including delta (p=0.21), theta (p=0.74), sigma (p=0.65), beta (p=0.85), or gamma (p=0.70). p<0.05 *, p<0.01 **. ns: not significant.
