## Supplemental Figure 6 for "Neuronal PAS domain 1 identifies a major subpopulation of wakefulness-promoting GABAergic neurons in basal forebrain"

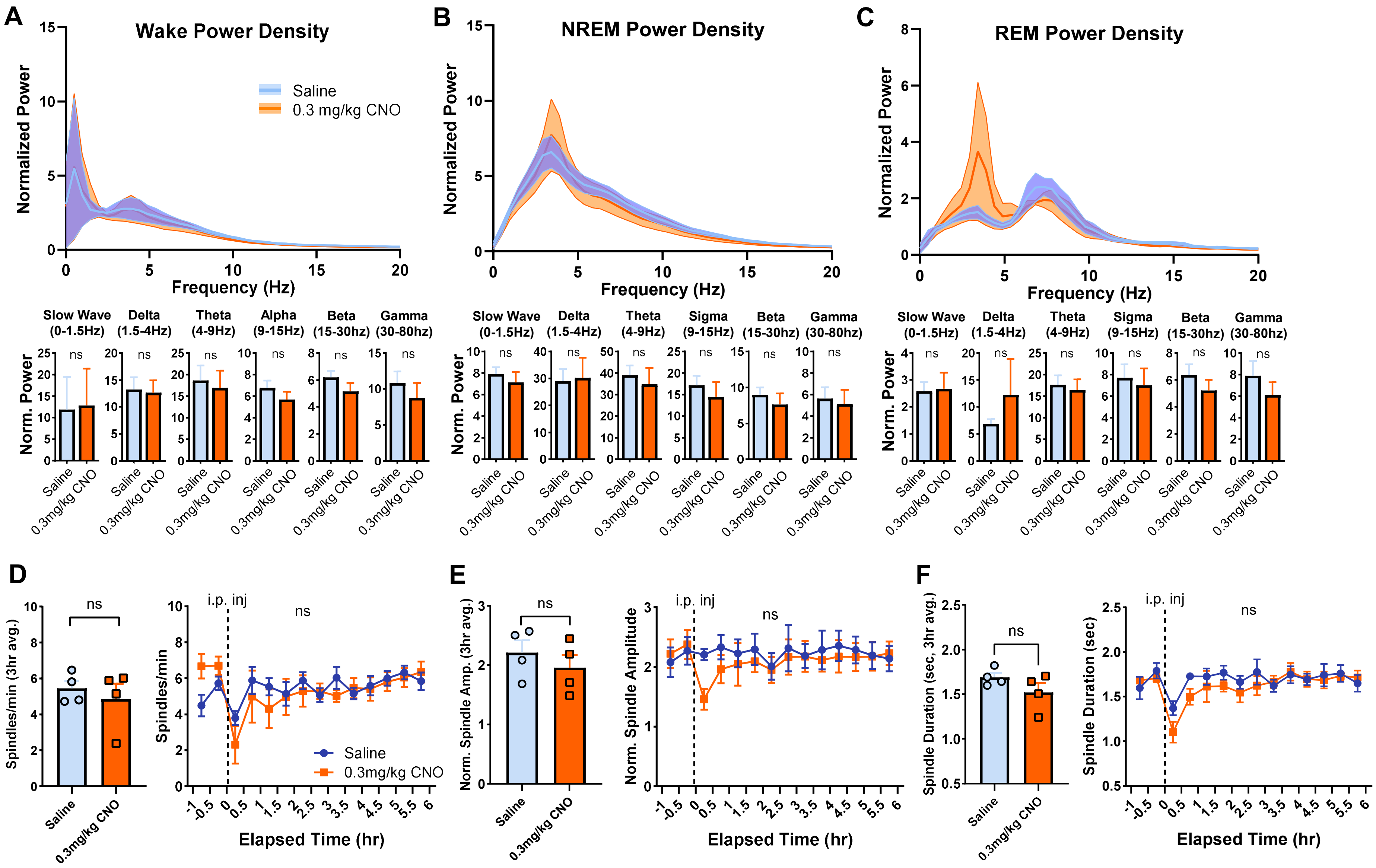
A
Wake Power Density
Normalized Power
Frequency (Hz)
Saline
0.3 mg/kg CNO
B
NREM Power Density
C
REM Power Density
Slow Wave (0-1.5Hz)
Delta (1.5-4Hz)
Theta (4-9Hz)
Alpha (9-15Hz)
Sigma (9-15Hz)
Beta (15-30hz)
Gamma (30-80hz)
Norm. Power
ns
Saline
0.3mg/kg CNO
D
Spindles/min (3hr avg.)
Spindles/min
i.p. inj
Elapsed Time (hr)
E
Norm. Spindle Amp. (3hr avg.)
Norm. Spindle Amplitude
F
Spindle Duration (sec, 3hr avg.)
Spindle Duration (sec)

**Supplemental Figure 6: CNO alone has no impact on cortical electroencephalogram (EEG) or sleep spindle activity. (A)** Without transduction of excitatory hM3D(Gq), administration of 0.3 mg/kg CNO (n=4) does not significantly change wake EEG power spectra compared to saline (n=4) across slow wave (0-1.5Hz, p=0.43), delta (1.5-4Hz, p=0.73), theta (4-9Hz, p=0.21), alpha (9-15Hz, p=0.19), beta (15-30Hz, p=0.14) or gamma bands (30-80Hz, p=0.31). Comparisons were made using two-tailed paired t-tests. **(B)** 0.3 mg/kg CNO alone does not significantly impact NREM EEG power spectra compared to saline at slow wave (p=0.33), delta (p=0.74), theta (p=0.27), sigma (p=0.13), beta (p=0.13), or gamma power bands (p=0.23). **(C)** REM power spectra is not significantly impacted by 0.3 mg/kg CNO vs. saline administration at slow wave (p=0.89), delta (p=0.43), theta (p=0.73), sigma (p=0.57), beta (p=0.18), or gamma bands (p=0.16). **(D)** Over the 3-hour post-injection period, spindle density was not significantly different between saline (5.45 ± 0.42 spindle/min) or 0.3 mg/kg CNO (4.86 ± 0.84 spindle/min, two-tailed paired t-test p=0.40). There is no significant difference in spindle density between treatments at any timepoint following injection. Two-way ANOVA identifies no interaction of time x treatment ($F_{13,74}$ = 1.53, p=0.13). **(E)** Normalized spindle amplitude did not significantly differ between saline (2.21 ± 0.21) and 0.3 mg/kg CNO (1.96 ± 0.22, p=0.16). Spindle amplitude did not significantly differ at any point following injection, and two-way ANOVA identifies no interaction of time x treatment ($F_{13,74}$ = 1.77, p=0.09). **(F)** Additionally, spindle duration following saline administration (1.69 ± 0.049 sec) was not significantly different than 0.3 mg/kg CNO treatment (1.52 ± 0.10 sec, p=0.21). Spindle duration was not significantly different at timepoints following drug injection, and two-way ANOVA identifies no interaction of time x treatment ($F_{13,74}$ = 1.19, p=0.30). ns: not significant.
