## Supplementary Table 1 for "Neuronal PAS domain 1 identifies a major subpopulation of wakefulness-promoting GABAergic neurons in basal forebrain"

| Antigen | Immunizing Specificity | Host Species | Source/ Catalog Number | Dilution | References (Mouse Brain Tissue) | Antibody Registry # |
|---|---|---|---|---|---|---|
| *Choline acetyltransferase (ChAT)* | 68-kDa band on western blot assays Source: Manufacturer; Bruce et al., 1995 | Goat | EMD Millipore AB144P | 1:200 | Sanz-Diez et al., 2019 Grady et al., 2020 | AB_2079751 |
| *Proto-oncogene c-Fos* | ~60/56 kDa band observed in Western Blot assay. Source: Manufacturer | Rabbit | EMD Millipore ABE457 | 1:200 | Yuan et al., 2017 Takata et al., 2018 | AB_2631318 |
| *Green Fluorescent Protein (GFP)* | 27-kDa monomer of 238 amino acids No detectable cross-reactivity with RFP Source: Manufacturer | Mouse | EMD Millipore MAB3580 | 1:300 | Helgager et al., 2013 Jo et al., 2018 Thankachan et al., 2019 | AB_94936 |
| *Neuronal PAS domain 1 (NPAS1)* | Validated in Npas1-cre-tomato and NPAS1 knockout mice (Hernandez et al., 2015) | Guinea Pig | Savio Chan (Northwestern Univ) | 1:1000 | Hernandez et al., 2015 Abecassis et al., 2020 | N/A |
| *Parvalbumin (PV)* | 12-kDa band on western blot assays Source: Manufacturer | Sheep | RnD AF5058 | 1:150 | Iijima et al., 2014 Usoskin et al., 2015 Stephany et al., 2016 | AB_2173907 |
| *Red Fluorescent Protein (RFP)/DSRed* | 30-38 kDa band on western blot assays Source: Manufacturer | Rabbit | Takara 632496 | 1:1000 | Pardo-Garcia et al., 2019 Grady et al., 2020 | AB_10013483 |

Supplementary table 1: primary antibodies.
