## Supplementary Table 2 for "Neuronal PAS domain 1 identifies a major subpopulation of wakefulness-promoting GABAergic neurons in basal forebrain"

| Secondary antibody | Vendor | Host Species | Catalog Number | Dilution | Incubation |
|---|---|---|---|---|---|
| *Donkey anti-mouse IgG AF488* | Thermo Fisher | Donkey | A21202 | 1:500 | 16 hrs 4ºC |
| *Donkey anti-goat IgG AF350* | Thermo Fisher | Donkey | A21081 | 1:100 | 4 hrs RT |
| *Donkey anti-goat IgG AF488* | Thermo Fisher | Donkey | A11015 | 1:100 | 4 hrs RT |
| *Donkey anti-guinea pig IgG AF488* | Jackson | Donkey | 706-545-148 | 1:500 | 2 hrs RT |
| *Donkey anti-guinea pig IgG AF594* | Jackson | Donkey | 706-585-148 | 1:500 | 2 hrs RT |
| *Donkey anti-rabbit IgG AF488* | Thermo Fisher | Donkey | A21206 | 1:200 | 2 hrs RT |
| *Donkey anti-rabbit IgG AF594* | Thermo Fisher | Donkey | A21207 | 1:100 | 2 hrs RT |
| *Donkey anti-sheep IgG AF350* | Thermo Fisher | Donkey | A21297 | 1:200 | 3 hrs RT |
| *Donkey anti-sheep IgG AF488* | Thermo Fisher | Donkey | A11015 | 1:200 | 3 hrs RT |

Supplemental table 2. Secondary antibodies
